# Molecular mechanism for the functional divergence between cohesin paralogs during meiosis

**DOI:** 10.64898/2026.05.26.727825

**Authors:** Purva, Ananya Dodamani, Mridula Nambiar

## Abstract

The key features of meiosis that enhance evolutionary success compared to mitosis are the processes of recombination and independent assortment during chromosome segregation, which help cells adapt to changing environments. Cohesins play critical roles in both these processes and have evolved specialized paralogs that are essential for meiotic chromosome dynamics. Studies so far have elucidated the roles of these meiotic cohesins but it is still unclear why the original mitotic proteins could not serve these evolved functions. In this study, we identify the mechanistic steps that are lost during meiosis when the mitotic counterparts replace the meiotic cohesins in *Schizosaccharomyces pombe*. The meiotic cohesin subunit Rec8^REC8^ has evolved multiple unique features that differentiate it from its mitotic paralog Rad21^RAD21^. Although ectopic expression of Rad21 in meiosis allows its chromatin enrichment, it fails to support reductional separation of chromosomes, initiation of recombination and protection of cohesion in anaphase I, resulting in catastrophic segregation errors. In contrast, the meiotic cohesin regulatory subunit Rec11^STAG3^, has only one major function of initiation of recombination, which is expectedly hampered in its absence. We show that although the mitotic paralog Psc3^STAG1/2^ is highly enriched at the cohesin-rich chromosomal axes, it is unable to recruit the downstream activators required for the induction of double-strand breaks. Our work systematically demonstrates the minimal functions that were necessary for the molecular evolution of these paralogs and explains the mechanisms that led to these adaptations.

## Introduction

Meiosis is a unique form of cell division that requires extensive remodeling of chromosome structures to support exchange of genetic information via recombination and achieve reduction in chromosome numbers ^1^. In *Schizosachharomyces pombe*, the hetero-tetrameric cohesin complex plays a pivotal role in initiation of meiotic recombination during prophase I that leads to generation of chiasma, which hold the homologous chromosomes together to facilitate proper chromosomal segregation in meiosis I (MI) ^2,3^. Furthermore, cohesins enriched at the centromeres and the flanking pericentromeric regions are equally critical to ensure accurate mono-orientation of sister kinetochores during MI and further equational segregation of sister-chromatids at meiosis II (MII) ^4^. The cohesin ring is made up of four core structural proteins, the SMC proteins, Psm1^SMC1^ (Human orthologs in superscript) and Psm3^SMC3^, the kleisin subunit Rad21^RAD21^ and a HEAT-repeat protein associated with kleisin (HAWK) Psc3^STAG1/STAG2^ (Fig. 1A) ^2^. The composition of the cohesin complex changes in cells undergoing meiosis, with mainly the Rad21^RAD21^ protein being replaced by Rec8^REC8^ and presence of Rec11^STAG3^ along with Psc3^STAG1/STAG2^, in fission yeast ^5^ (Fig. 1A and S1). Rec8^REC8/RAD21L^ and Rec11^STAG3^ are expressed only in meiotic cells or in mammalian germ cells. Interestingly in mammalian cells SMC1 protein has two paralogs SMC1α and SMC1β, with the latter exclusively expressed in the germ cells ^6^.

**Figure 1.**
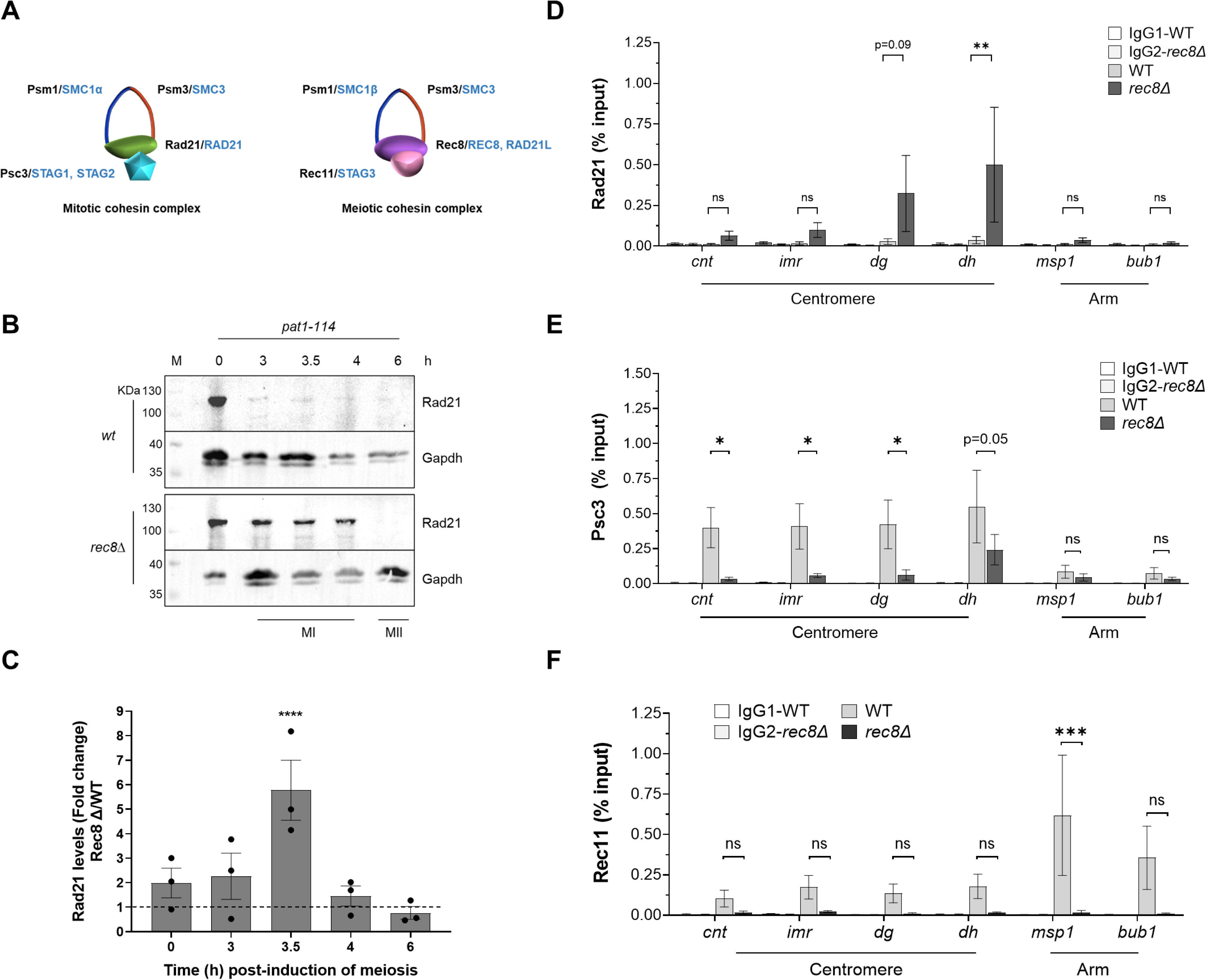
Rad21 expression and chromatin enrichment increases at the centromeres in *rec8Δ.* **A.** Schematic representation of cohesin complexes present in *S. pombe* (black) and *H. sapiens* (blue) during mitosis and meiosis. **B.** Western blot showing expression levels of Rad21 during *rec8+* and *rec8Δ* upon induction of meiosis in *a pat1-114* strain. The blots were probed with anti-Rad21 and anti-Gapdh antibodies. 3-4 h post-induction marks MI stage and between 6-7 h constitutes the MII stage during *pat1-114* induced meiosis. M is the molecular weight ladder and the sizes are marked. **C.** Quantification of Rad21 expression in *rec8Δ* expressed as fold change relative to that in wild type. **D – F.** ChIP qPCR for Rad21 (D), Psc3 (E) and Rec11 (F) in *wt* and *rec8Δ* at centromere (*cnt, imr, dg* and *dh*) and arm (*msp1* and *bub1*) loci. Pulldowns with IgG are used as controls. The bars represent the mean of at least three independent experiments performed in triplicates; error bars=SEM. ****p <0.0001; ***p <0.001; **p<0.01; *p <0.05; ns p >0.05. The data were analysed using Two-way ANOVA test (Tukey’s multiple comparisons test).

Meiotic kleisin Rec8 partnered with the meiotic HAWK protein Rec11 have been shown to be necessary for initiation of meiotic DNA double-strand breaks (DSBs) induced by Rec12^SPO11^ ^7,8^. Meiotic chromosomes are organized into higher-order structures composed of axes and loops, primarily established by the meiotic Rec8-Rec11 cohesins ^9^. The DSB-inducing Rec12-Rec14-Rec6 complex (DSBC) sits on the chromosomal axis and gets activated by the linear element (LinE) protein Rec10, which is recruited to the axis upon phosphorylation of Rec11 by the Hhp1-Hhp2 (CK1) kinase complex ^10,11^ (Fig. S2A). DSB hotspots are marked by the other LinE complex proteins Rec25-Rec27-Mug20 and are present on the loops, which are brought close to the axis by the mediator Mde2 that interacts with the Rec7-Rec15-Rec24 (SFT) complex at the hotspot ^8,12^. All these interactions converge to form the tethered loop-axis complex, which is critical for the activation of DSBs and facilitates regulation of DSB numbers and recombination patterning on the chromosomes ^9,13–15^.

The presence of distinct cohesin complexes in meiosis argues for vital functions that have evolved specifically in meiotic cells and may not be complemented by the mitotic paralogs. Indeed, yeast strains that are deficient in either Rec8 or Rec11 have provided insights into their indispensable roles in meiosis-specific processes ^7,8,16^. In mice, the mitotic SMC1α could substitute for most of the meiotic functions of SMC1β, except for rescue of telomere damage ^17^. Hence, the degree of specialization adopted by meiotic paralogs and the similarities with their mitotic counterparts can vary for the protein pair, across species. Moreover, several questions remain regarding how this functional divergence evolved, how is it maintained mechanistically and the extent of the overlapping and unique functions of these paralogs. Such systematic analyses are critical to answer why such new evolved functions were necessary during evolution of sexual reproduction.

In this study, we systematically dissect the functional redundancies and non-overlapping functions of the two meiosis-specific cohesins, the Kleisin Rec8^REC8^ and HAWK protein Rec11^STAG3^ with their mitotic paralogs, Rad21^RAD21^ and Psc3^STAG1/2^, respectively, in fission yeast meiosis. We find that mitotic subunits can form a functional complex in the absence of their respective meiotic paralogs and do get loaded on the chromosomes, but fail to perform specific functions such as initiation of meiotic recombination and proper reductional division. Our data shows that the meiotic paralogs have gained specialized functions via new critical interactions that their mitotic counterparts lack and we demonstrate how the absence of these downstream steps affect the critical meiosis-specific functions. This stringent exclusivity of functions stems from the otherwise catastrophic repercussions that these meiotic features may have during mitotic division. Our work is a methodical analysis that provides fundamental insights to answer why meiosis-specific paralogs have emerged during evolution and the molecular basis of how their distinct functions are maintained.

## Results

### Native Rad21 cannot complement functions in the absence of Rec8 despite chromosomal occupancy at MI

Previous studies have shown that in the absence of Rec8 both meiotic recombination and segregation functions are severely compromised ^7,8,16^. In terms of structures, both Rec8 and its mitotic paralog Rad21 exhibit significant similarities at the highly conserved N and C-terminal domains, while a third middle domain that is proposed to bind with the HAWK subunits, shows differences as seen in the Alpha Fold predicted structures ^18,19^ (Fig. S1A). The native expression of Rad21 is reduced upon induction of meiosis in wild type, as is expected, allowing Rec8 to be effectively the only kleisin for cohesin ring formation (Figs. 1B, S2A and S2B). Interestingly, in *rec8Δ* cells, the native Rad21 levels persist throughout prophase I (4 hpi), where recombination predominantly occurs (Figs. 1B, 1C). We checked if this Rad21 population shows any chromosomal occupancy during prophase I, and found significant chromatin-bound fraction at the outer repeats (*dh*) at the pericentromeric regions as compared to wild type where Rad21 expression is absent (Fig. 1D). There was no occupancy at the chromosomal arms (*msp1, bub1*) (Fig. 1D). To see if other partner cohesin subunits also follow similar chromatin binding patterns, we checked for the two potential HAWK partners, Psc3 and Rec11, and found that as expected levels of Psc3 dropped significantly in *rec8Δ* specifically at the centromeric loci (*cnt, imr, dg, dh*), while Rec11 reduced majorly at the chromosomal arms (*msp1-* cohesin rich locus) (Figs. 1E, 1F). Psc3 did retain some occupancy at the outer centromeric repeats (*dh*), suggesting minimal loading of the Rad21-Psc3 complex (Fig. 1E and S2D). Hence, in the absence of Rec8, there appears to be minimal cohesin loading atleast at the pericentromeric repeats. However, this is not sufficient to maintain proper chromosomal segregation, which was confirmed when we used a fluorescence-based tetrad assay, described previously ^20,21^. We used this assay to classify the segregation defects into major stage-specific errors (MI and MII) as well as sub-classified them based on mechanism such as homolog non-disjunction (NDJ), premature separation of sister chromatids (PSSC) or reverse segregation (RS; equational segregation) (Figs. 2A and S3). Our assay nicely captures all these errors in *rec8Δ* tetrads, which show predominantly MI PSSC errors due to loss of pericentric cohesion, MI NDJ due to loss of meiotic recombination and hence random separation of homologs, as well as reverse segregation due to impaired mono-orientation of sister-kinetochores in the absence of cohesion at the centromere core (Fig. 2B and S4). Since Rad21 protein levels are near absent at MII stage in *rec8Δ*, one would expect to see increased mis-segregation at MII due to loss of sister-chromatid cohesion (Fig. 1B, 1C). We indeed found that nearly all the tetrads with errors originating at MI, further showed MII defects (Figs. 2C, 2D). Hence, with respect to cohesion atleast, it appears that perhaps lack of sufficient Rad21 through to the MII stage may be a reason for no rescue of cohesion defects.

**Figure 2.**
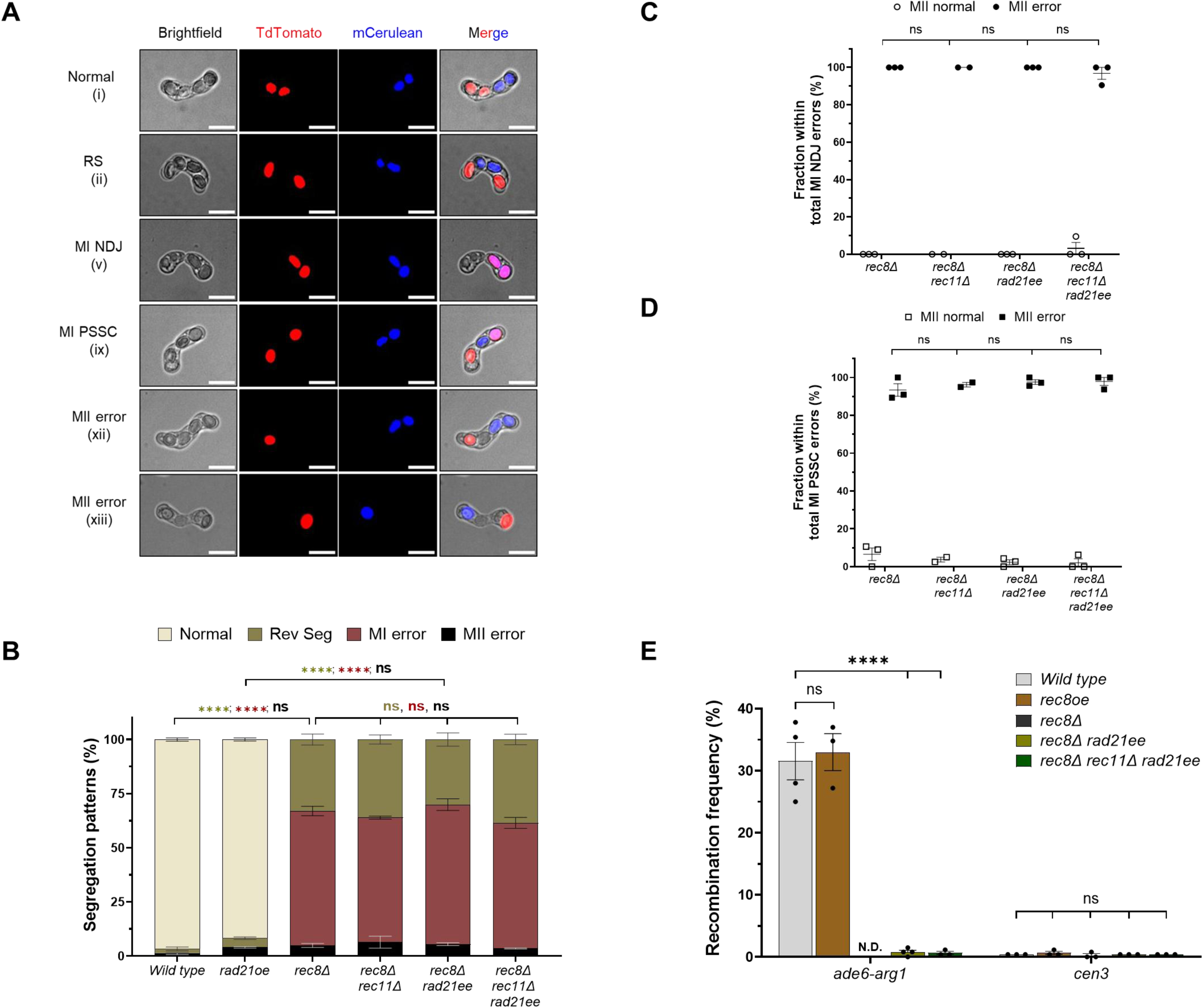
Overexpression of Rad21 fails to restore the segregation and recombination functions in *rec8Δ* meiosis. **A.** Representative images for normal, reverse segregation (RS), meiosis I non-disjunction (MI NDJ), meiosis I premature separation of sister chromatids (MI PSSC) and meiosis II (MII) error types. All the examples of tetrads as classified in Fig. S3A are shown along with the corresponding numbers as depicted in the schematic cartoon. Scale bars represent 10 μm. **B.** Plot showing frequency of normal, RS, MI and MII segregation errors in wild type*, rad21oe, rec8Δ, rec8Δ rec11Δ, rec8Δ rad21ee* and *rec8Δ rec11Δ rad21ee* genotypes. **C-D.** Graph showing fraction of normal and erroneous MII segregation within MI NDJ (C) and MI PSSC (D) events in *rec8Δ, rec8Δ rec11Δ, rec8Δ rad21ee* and *rec8Δ rec11Δ rad21ee* genotypes. All data are mean ± SEM (n ≥ 3 experiments; 2 for *rec8Δ rec11Δ*), assaying 563, 506, 362, 208, 438 and 395 tetrads for wild type*, rad21oe, rec8Δ, rec8Δ rec11Δ, rec8Δ rad21ee* and *rec8Δ rec11Δ rad21ee*, respectively. **E.** Recombinant frequency (RF) across *ade6-arg1* arm interval and *cen3* in wild type*, rec8oe, rec8Δ rad21ee* and *rec8Δ rec11Δ rad21ee* genotypes. Data are mean ± SEM (n ≥ 3 experiments), assaying 509, 396, 527 and 396 spore colonies for wild type*, rec8oe, rec8Δ rad21ee* and *rec8Δ rec11Δ rad21ee,* respectively, for arm recombination and 395, 396, 371, 396 and 395 spore colonies for *cen3* recombination in wild type*, rec8oe, rec8Δ, rec8Δ rad21ee* and *rec8Δ rec11Δ rad21ee,* respectively. ****p <0.000; ns p >0.05. Data in panels B-D were analysed by Two-way ANOVA test (Tukey’s multiple comparisons test), whereas Two-tailed Fisher’s exact test was used for data in panel E. The data for individual experiments for panels B-E are provided in Tables S3, S5 and S6.

### Ectopic Rad21 expression does not restore segregation fidelity during *rec8Δ* meiosis

Unavailability of Rad21 at MII could be due to either lower expression compared to Rec8 or lack of protection of the pericentromeric population at anaphase I. To test the former, we ectopically expressed Rad21 from *rec8* promoter (denoted as *rad21ee*) to match the native level of kleisin available in *rec8Δ* cells compared to wild type. However, *rad21ee* still did not improve reductional segregation at MI as seen by similarly high levels of RS tetrads nor did it alleviate any of the MI errors seen in *rec8Δ* or *rec8Δ rec11Δ* (Fig. 2B and S4). Subsequent MII errors in the tetrads having defects originating in MI such as NDJ or PSSC, were unchanged as compared to *rec8Δ* (Fig. 2C, 2D and S4C). This clearly indicates that increasing Rad21 levels upto that of native Rec8 is not sufficient to restore the specialized functions of the meiotic paralog. Although, Rad21 and Rec8 are similar with respect to their ability to form a complex with Psc3 and get loaded at the centromeres, lack of Rad21-Psc3 mediated cohesion at the centromeric core in *rec8Δ* as compared to wild type explains equational segregation and biorientation of sister kinetochores at MI (Fig. 1D, 1E, 2B, S2C and S2E). Additionally it is also possible that the Rad21-Psc3 at the outer pericentromeric regions may not be protected like Rec8 during anaphase I, thereby leading to elevated MII errors ^4^. Hence, evolution of Rec8’s cohesion function helped in solving the way to perform reductional division of homologs and maintain sister-chromatid cohesion till MII.

### Improper linear element formation in *rec8Δ* despite forced Rad21-Rec11 association leads to recombination defects during MI

Next, we measured recombination at both the chromosomal arms (*ade6-arg1* interval) and centromeres (*cen3)* in the presence of *rad21ee* and observed no changes in recombination frequency levels compared to *rec8Δ* (Fig. 2E). Centromeric recombination remained repressed in all the genotypes tested, whereas, arm recombination was totally abolished in *rec8Δ* when compared to wild type. We also removed Rec11 to check if having only one functional Rad21-Psc3, without interference from the meiotic HAWK has any effect on recombination, and found no difference (Fig. 2E).

To elucidate the reasons behind why Rad21 does not support recombination, despite being sufficiently expressed during MI, we wondered if there were defects in the steps downstream to cohesins in the recombination pathway (Fig. 1B, 1C and S2A). Since phosphorylation of Rec11 is critical for recruiting the linear element complex containing Rec10 that is essential to activate the Spo11 complex, we used both a *rec11-5D* phosphomimic and a *rec11-rec10* fusion allele that are sufficient to rescue *hhp1-hhp2* mutants and *rec10Δ* for recombination, as described earlier ^10^. *rad21ee rec11-5D* or *rec11-rec10* in *rec8Δ* should bypass the potential defect for recombination, if phosphorylation of the Rad21-Rec11 complex and subsequent recruitment of Rec10 are the limiting steps. However, we did not observe any major rescue in the levels of arm recombination nor improvement in the recombination-associated segregation defects in these genotypes (Fig. 3A, 3B). We did notice a small but significant increase in only one class of recombinants in the *rec11-rec10* allele, probably indicating that these may not be arising from reciprocal crossover events and may involve gene conversions. However, it was still significantly low as compared to the wild type recombination levels (Fig. 3A). Additionally, *rec11-rec10* couldn’t alleviate any of the MI errors in the same background and the overall frequencies of the segregation defects were similar, confirming lack of rescue (Fig. 3B).

**Figure 3.**
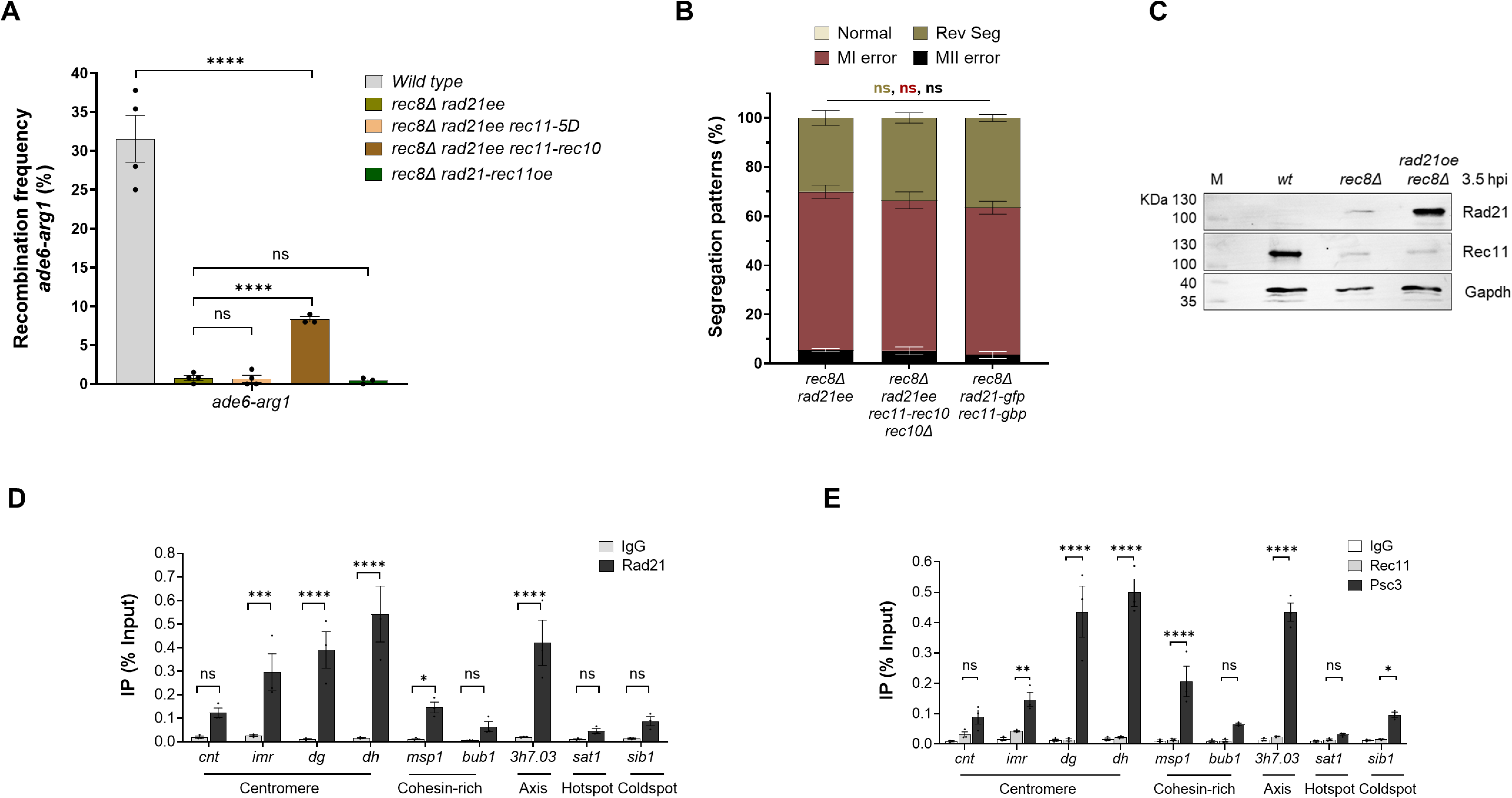
Absence of robust Rad21-Rec11 cohesin complex during *rec8Δ* meiosis leads to recombination and segregation defects. **A.** Recombinant frequency (RF) across *ade6-arg1* arm interval in wild type*, rec8Δ rad21ee, rec8Δ rad21ee rec11-5D, rec8Δ rad21ee rec11-rec10* and *rec8Δ rad21-rec11oe* genotypes. Data are mean ± SEM (n ≥ 3 experiments), assaying 509, 527, 406, 567 and 393 spore colonies for wild type*, rec8Δ rad21ee, rec8Δ rad21ee rec11-5D, rec8Δ rad21ee rec11-rec10 rec10Δ* and *rec8Δ rad21-rec11oe,* respectively. **B.** Plot showing frequency of normal, RS, MI and MII segregation errors in *rec8Δ rad21ee, rec8Δ rad21ee rec11-rec10 rec10Δ* and *rec8Δ rad21-gfp rec11-gbp* genotypes. All data are mean ± SEM (n ≥ 3 experiments), assaying 438, 373 and 316 tetrads for *rec8Δ rad21ee, rec8Δ rad21ee rec11-rec10 rec10Δ* and *rec8Δ rad21-gfp rec11-gbp*, respectively. **C.** Western blot showing expression levels of Rad21 and Rec11 in wild type, *rec8Δ* and *rec8Δ rad21oe* at 3.5 h post-induction in *a pat1-114* meiosis. 3-4 h post-induction marks MI stage and between 6-7 h constitutes the MII stage during *pat1-114* induced meiosis. The blots were probed with anti-Rad21, anti-HA (for Rec11-HA) and anti-Gapdh antibodies. M is the molecular weight ladder and the sizes are marked. **D-E.** ChIP qPCR for Rad21 (D), Psc3 (E) and Rec11 (E) in *rec8Δ rad21oe* genotype at centromere (*cnt, imr, dg* and *dh*) and various arm loci such as cohesin-rich (*msp1, bub1*), LinE/Axis (*3h7.03),* DSB hotspot (*sat1*) and coldspot (*sib1)*. Pulldowns with IgG were used as controls. The bars represent the mean of at least three independent experiments performed in triplicates; error bars = SEM. For each experiment, atleast 3 biological repeats were performed. ****p <0.0001; ***p <0.001; **p<0.01; *p <0.05; ns p >0.05. Data in panel A was analysed by Two-tailed Fisher’s exact test, whereas Two-way ANOVA test (Tukey’s multiple comparisons test) was used for data in panel B. The data for individual experiments in panels A and B are provided in Tables S3 and S5, respectively.

Although Rad21-Rec11 complex formation is not limiting, as seen previously, it doesn’t appear to be as robust as Rad21-Psc3 ^5,18^. For confirming the Rad21-Rec11 interaction during meiosis I, we overexpressed Rad21 under the *Padh1* promoter (*rad21oe*), and first checked for the expression of both Rad21 and Rec11 in the chromatin fraction. To our surprise, we observed near absent expression of Rec11 in both *rec8Δ* and *rec8Δ rad21oe* as compared to wild type (Fig. 3C). Rad21 was clearly present in *rec8Δ* and overexpressed in *rec8Δ rad21oe* cells (Fig. 3C). We further confirmed the chromatin occupancy of the overexpressed Rad21 at the chromosomal arms and pericentromeric loci, but not in the centromeric core (Fig. 3D). The chromatin-bound levels of the overexpressed Rad21 were quite similar to that observed in *rec8Δ* with wild type Rad21 (Fig. 1D, 3D). There was no enrichment of Rec11, as expected and Psc3 was present at the pericentromeric loci along with Rad21 (Fig. 3E). We also saw increased occupancy of Rad21-Psc3 complex at the cohesin-rich chromosomal axis regions (*msp1* and *3h7.03*) (Fig. 3D, 3E). This indicates that in the absence of Rec8, its meiotic partner Rec11 may not be as stable as in wild type and consequently has a lowered expression.

In order to bypass the lowered expression of Rec11 and forcing its complex formation with Rad21, we employed two approaches. In the first, we directly fused Rec11 to the C-terminus of Rad21 (*rec8Δ::Padh1-rad21-rec11* allele) (Fig. S5A). We confirmed its functionality by expressing it in the temperature sensitive *rad21-k1* cells and observed rescue of growth at the restrictive 34°C (Fig. S5B, S5C). However, the fusion allele did not rescue recombination (Fig. 3A). Our second approach was using a GFP-binding protein (GBP) – GFP system, wherein Rec11 was expressed under a constitutive promoter and fused with GBP, such that it forcefully interacts with Rad21-GFP (Fig. S5D). Again, we did not observe any reduction in *rec8Δ* associated segregation errors (Fig. 3B). Despite the overexpression of Rec11, these strains still showed recombination defects, indicating that formation of stable chromatin-bound Rad21-Rec11 complex may be the major limiting step when Rec8 is absent in wild type cells. Moreover, the mitotic Rad21-Psc3 complex, although enriched at the cohesin-rich axes at the chromosomal arms in *rec8Δ* fail to activate recombination, suggesting functional divergence between the HAWK subunits Psc3 and Rec11 as well.

### Mitotic HAWK subunit Psc3 and meiotic Rec11 have mutually exclusive functions

We next aimed to understand the divergence between the HAWK subunits Psc3 and Rec11. In terms of Alpha Fold structures, both these proteins exhibit similarities at the conserved STAG and Stromalin domains, whereas the HEAT repeat containing domain that is crucial for its interaction with other proteins, appears to be dissimilar in terms of sequence and structure level ^19^ (Fig. S1A). We initially confirmed the differential enrichment of the mitotic Psc3 at the centromeric loci, while the meiotic Rec11 was enriched at the chromosomal arms ^5,10^ (Fig. S2E). Both *rec11Δ* and *rec11-5A* phosphodeficient mutant that fails to get phosphorylated by *hhp1-hhp2* are strongly defective in chromosomal arm recombination as tested at the genetic interval *ade6-arg1* (Fig. 4A). On the contrary, *psc3Δ* that requires concomitant ectopic expression of Rec8 and Rec11 for mitotic cell viability, has no significant effect on recombination frequency in the arms (Fig. 4A). None of the strains tested showed any derepression of recombination at the centromeres and were identical to the wild type.

**Figure 4.**
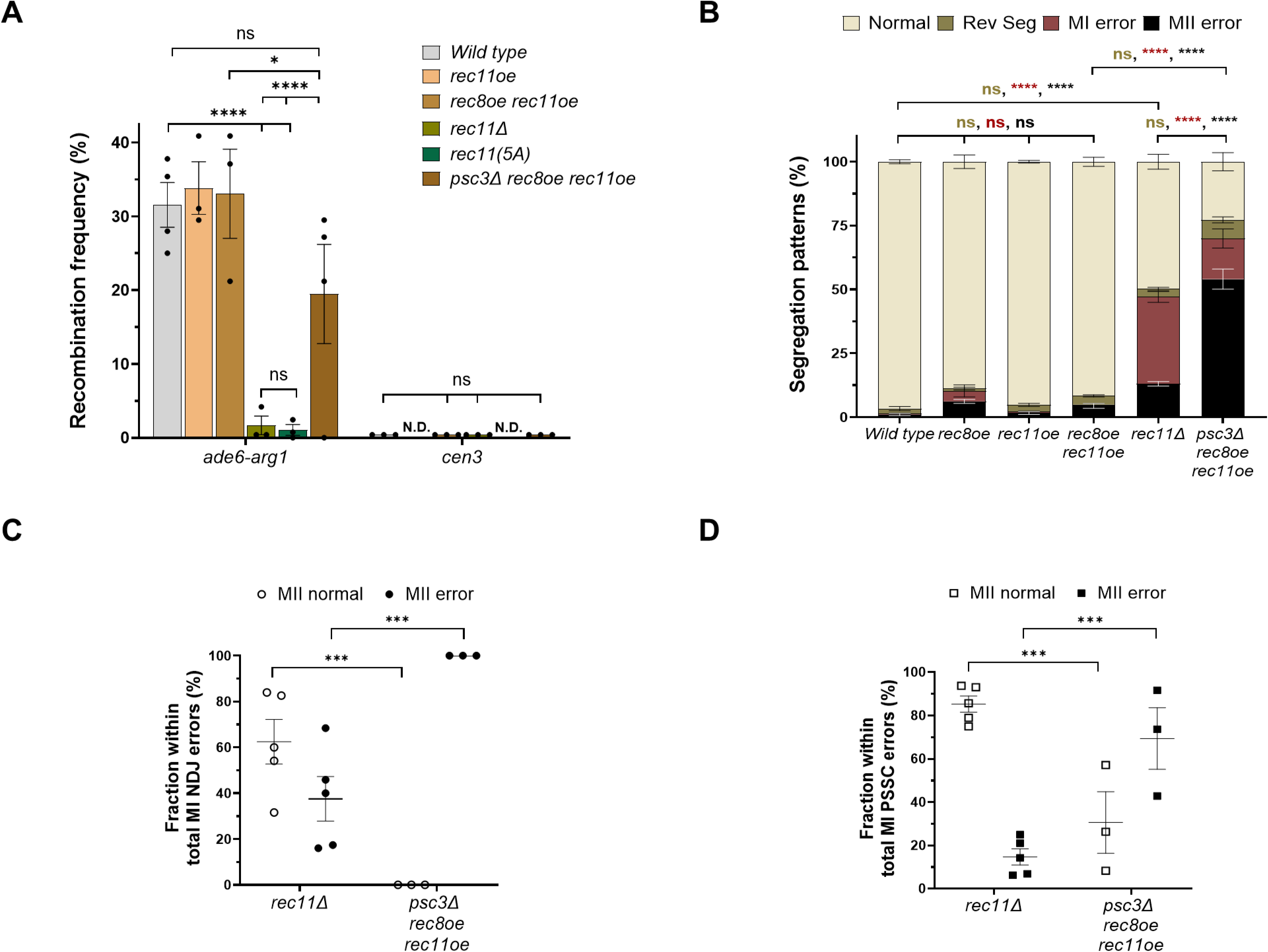
The two HAWK paralogs have distinct functions during recombination and segregation in meiosis. **A.** Recombinant frequency (RF) across *ade6-arg1* arm interval and *cen3* in wild type*, rec11oe, rec11oe rec8oe, rec11Δ, rec11(5A)* and *psc3Δ rec8oe rec11oe* genotypes. Data are mean ± SEM (n ≥ 3 experiments), assaying 509, 396, 396, 400, 386 and 396 spore colonies for wild type*, rec11oe, rec11oe rec8oe, rec11Δ, rec11(5A)* and *psc3Δ rec8oe rec11oe,* respectively. **B.** Plot showing frequency of normal, RS, MI and MII segregation errors in wild type, *rec8oe, rec11oe, rec8oe rec11oe, rec11Δ* and *psc3Δ rec8oe rec11oe* genotypes. **C-D.** Graph showing fraction of normal and erroneous MII segregation within MI NDJ (C) and MI PSSC (D) events in *rec11Δ* and *psc3Δ rec11oe rec8oe* genotypes. All data are mean ± SEM (n ≥ 3 experiments), assaying 563, 459, 659, 460, 614 and 289 tetrads for wild type, *rec8oe, rec11oe, rec8oe rec11oe, rec11Δ* and *psc3Δ rec8oe rec11oe*, respectively. ****p <0.0001; ***p <0.001; *p <0.05; ns p >0.05. Data in panel A was analysed by Two-tailed Fisher’s exact test, whereas Two-way ANOVA test (Tukey’s multiple comparisons test) was used for data in panels B-D. The data for panels A-D are provided in Tables S4, S5 and S6, respectively.

As expected, loss of recombination resulted in predominantly MI errors in *rec11Δ* showing near equal distribution of NDJ and PSSC errors, followed by significant MII errors and no discernible RS errors (Fig. 4B, S6). In contrast, *psc3Δ* showed a high ∼60% defect at MII stage and relatively lower errors at MI, when compared to *rec11Δ* (Fig. 4B, S6). Within MI errors, *psc3Δ* showed a higher propensity for PSSC events and very low NDJ errors, pointing towards loss of centromeric cohesion as the cause of segregation errors (Fig. S6). However, as was the case for *rec8Δ* above, the *psc3Δ* tetrads with an existing MI NDJ or PSSC were found to have a high frequency of subsequent MII errors as well, but this wasn’t the case for *rec11Δ* (Fig. 4C, 4D, 2C, 2D). High MII errors suggest lack of centromeric cohesion and indeed, we found that both Psc3 and Rec8 levels did not change across any of the loci tested either at the centromere or the arms, in *rec11Δ* compared to wild type (Fig. 5A, 5B). This explains the less frequent MII errors in *rec11Δ* and a higher frequency of normal MII segregation, in those cells that underwent either MI NDJ or PSSC events, indicating largely intact sister-chromatid centromeric cohesion (Fig. 4C, 4D-E).

**Figure 5.**
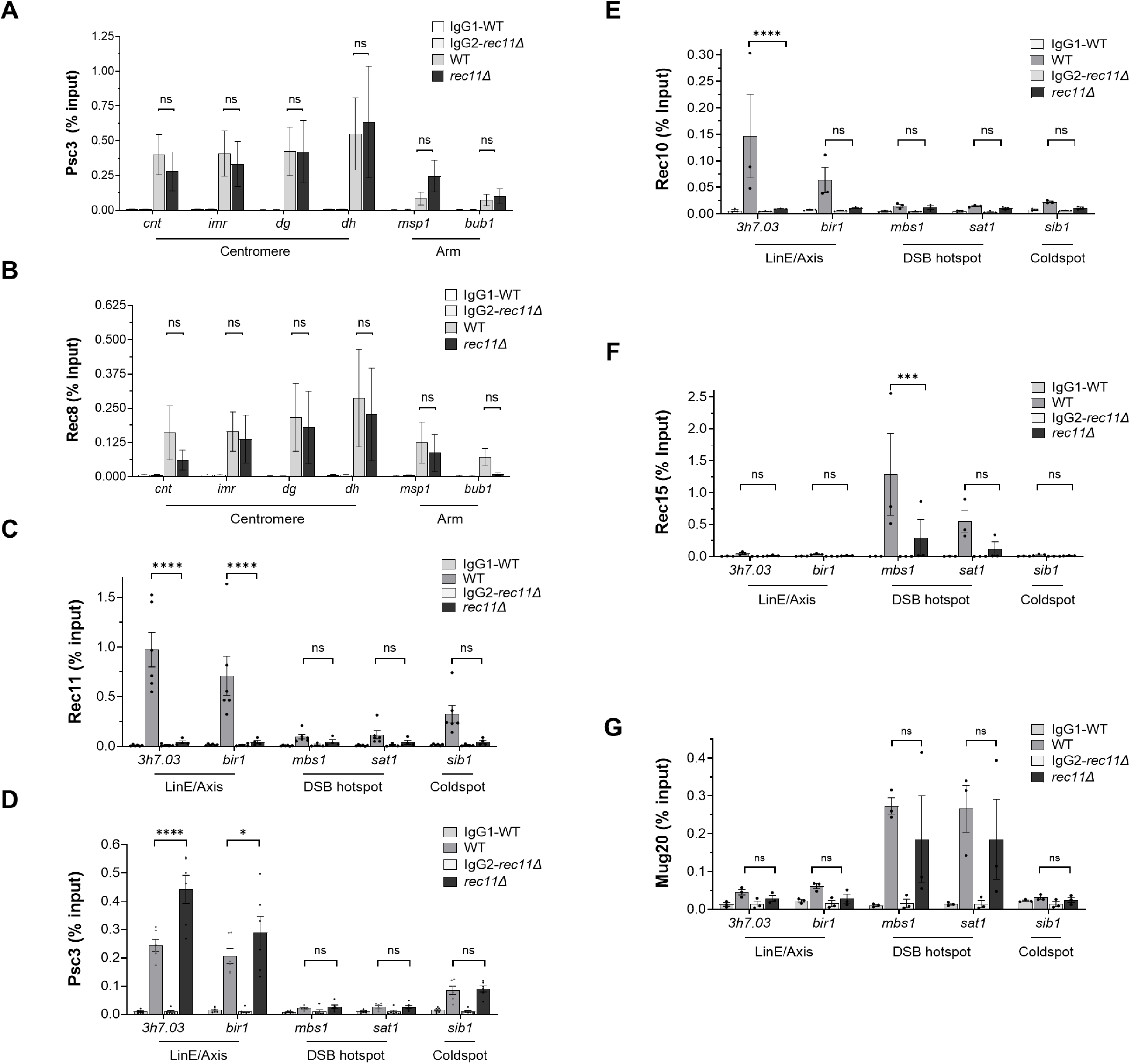
Psc3 containing cohesins are unable to recruit the DSB complex activator Rec10 to the chromosome axes unlike Rec11. A-B. ChIP qPCR for Psc3 (A) and Rec8 (B) in wild type and *rec11Δ* genotypes at centromere (*cnt, imr, dg* and *dh*) and arm (*msp1* and *bub1*) loci. **C-G.** ChIP qPCR for Rec11 (C), Psc3 (D), Rec10 (E), Rec15 (F) and Mug20 (G) in wild type and *rec11Δ* genotypes at various arm loci such as LinE/Axis (*3h7.03* and *bir1*), DSB hotspots (*mbs1* and *sat1*) and a coldspot (*sib1)*. Pulldowns with IgG were used as controls. The bars represent the mean of at least three independent experiments performed in triplicates; error bars=SEM. ****p <0.0001; ***p <0.001; *p <0.05; ns p >0.05. The data were analysed using Two-way ANOVA test (Tukey’s multiple comparisons test).

Overall, these data clearly show that Rec11 has evolved exclusively for activation of recombination, while Psc3 plays a stronger role with respect to cohesion function, especially at the centromeres. This also explains the logic behind spatial separation of chromatin occupancy for the two paralogs, with Rec11 excluded from near the centromeres, to restrict meiotic recombination events only to the chromosomal arms ^5,22^.

### Psc3 fails to support the tethered loop-axis complex for initiation of double-strand breaks

Since Psc3 did not support meiotic recombination, we wanted to understand the differences in its downstream molecular interactors such that only Rec11 can activate the Rec12 complex for initiating DSBs. We analysed known linear element/axis (*3h7.03* and *bir1*) loci that are known to be enriched for Rec8-Rec11, DSB hotspots (*mbs1* and *sat1*) and a coldspot (*sib1*) region, described earlier^10,12^. As expected, we found significant enrichment for Rec11 mainly at the LinE/axis loci, confirming the presence of the meiotic cohesins on the chromosomal axes (Fig. 5C). Interestingly, we found elevated levels of Psc3 on the LinE/axis loci in *rec11Δ,* clearly indicating that Rec8-Psc3 cohesins can compensate for the absence of Rec11 on the chromosomal axis (Fig. 5D). However, this was not followed by an enrichment of the main activator of Rec12 complex, Rec10, on this LinE/axis loci in *rec11Δ,* as seen in the wild type (Fig. 5E). As mentioned above, Rec15 from the SFT complex is mostly present on the DSB hotspots, which are primarily decorated by the LinE protein complex of Rec25-Rec27-Mug20 ^8,12^. We also observe enrichment of Rec15 and Mug20, specifically on the DSB hotspots in wild type, further providing evidence supporting the loop-axis model for DSB initiation (Figs. 5F, 5G, S7). In *rec11Δ,* there was a significant decrease in the chromatin enrichment of Rec15, suggesting loss of critical interactions between Rec10-Rec15 that help tether the hotspots to the chromosomal axis, thereby facilitating action of Rec12 containing DSB complex in DSB formation (Fig. 5F). As is expected, loss of Rec11 on the chromosome axis didn’t alter Mug20 enrichment on the hotspots present on the loops (Fig. 5G). Our data clearly demonstrate that despite Psc3 cohesin being able to contribute to chromosomal axis formation along with Rec8, it fails to recruit and interact with the downstream players in the pathway that are necessary to bring about the loop-axis tethering, which is the key in activating Rec12 complex to make the DNA breaks and initiate recombination.

## Discussion

It is largely believed that meiosis originated from mitosis in early eukaryotes and one of the selection pressures may possibly have been the need for genetic variation ^23^. Opposing views propose a primitive meiosis-like pathway that was utilized by early eukaryotes and mitosis to have been derived from it as a streamlined form of meiosis ^24^. Either way, the earliest unique step in meiosis may have been the extensive pairing and synapsis between homologous chromosomes that is foundational for recombination. Other distinctive properties of meiosis are the suppression of separation of sister chromatids, absence of replication in the second meiotic division and the protection of centromeric cohesion at the end of anaphase I to facilitate proper meiosis II segregation. It is intuitive that many of these specialized processes in meiosis may have evolved from the pre-existing mitotic machinery as many of the involved proteins have paralogs that came about post gene duplication events. The cohesin complexes are one such indispensable group of proteins that have essential functions in chromosome segregation and genome organization. Hence, it is not surprising that many of the cohesin structural subunits and their regulators have dedicated meiosis-specific paralogs that have evolved specialized functions. In this study, we aimed to systematically dissect out the extent of functional overlap and divergence among the kleisin and HAWK subunits of the cohesin complex. We find that mitotic proteins fail to perform the meiosis-specific functions of initiation of recombination and mono-orientation of homologs to prevent sister-chromatid separation. We also provide molecular evidence for the “missing” steps when mitotic paralogs are present that make them functionally inept in carrying out these pathways.

### Functional divergence between cohesin paralogs

Studies in *S. pombe* have elucidated the essential functions of the meiotic kleisin Rec8 and HAWK partner Rec11 in induction of meiotic DSBs as in their absence, recombination frequencies across genetic intervals are near abolished^7^. The discovery that both the HAWK subunits are expressed in meiosis, partner with Rec8 and have distinct spatial localization on the chromosomes, further led to the understanding that exclusion of Rec11 from the centromeres ensures absence of deleterious recombination and promote proper segregation during meiosis I ^5,22,25^. Forced enrichment of Rec11 complex to the centromeres increased recombination resulting in elevated mis-segregation events during meiosis I, pointing towards the inability of Psc3 to activate recombination ^20,22^. We provide evidence that recruitment of Rec10 and other DSB complex activation proteins, cannot happen when Psc3 is part of the cohesin complex on the chromosomal axis. N-terminal of Rec11 gets phosphorylated by CK1 kinase that triggers the recruitment of Rec10, which may be the limiting step when Psc3 is present ^10,11^. In absence of Rec8, Rad21 has been shown to occupy heterochromatin clusters during prophase I and we further confirm the enrichment of Rad21 at pericentromeric repeats ^4,26^. However, despite its presence, reductional segregation of homologous cannot happen as Rad21 doesn’t get deposited at the centromere core to facilitate monopolar attachment, unlike Rec8.

Proper meiosis II segregation relies on the protected cohesin population at the centromeres, which may be specific to centromeric Rec8, since we observe disappearance of Rad21 at the MII stage ^4,27^. In addition, ectopically increasing the Rad21 levels does result in robust cohesin loading at the cohesin-rich chromosomal axes as well, but still does not lead to activation of DSB formation. This shows that in the absence of Rec8, Rad21 is able to form a functional cohesin complex that can get loaded at the pericentromeres. However, since Psc3 is the only HAWK present in *rec8Δ*, due to lowered levels of Rec11, activation of recombination is still lacking. Hence, in fission yeast we propose that the major limitation for lack of complementation of recombination in absence of both the meiotic paralogs, is imposed due to the inability of Psc3 to interact with downstream DSB inducing complexes, irrespective of it partnering Rec8 or Rad21 (Fig. 6).

**Figure 6.**
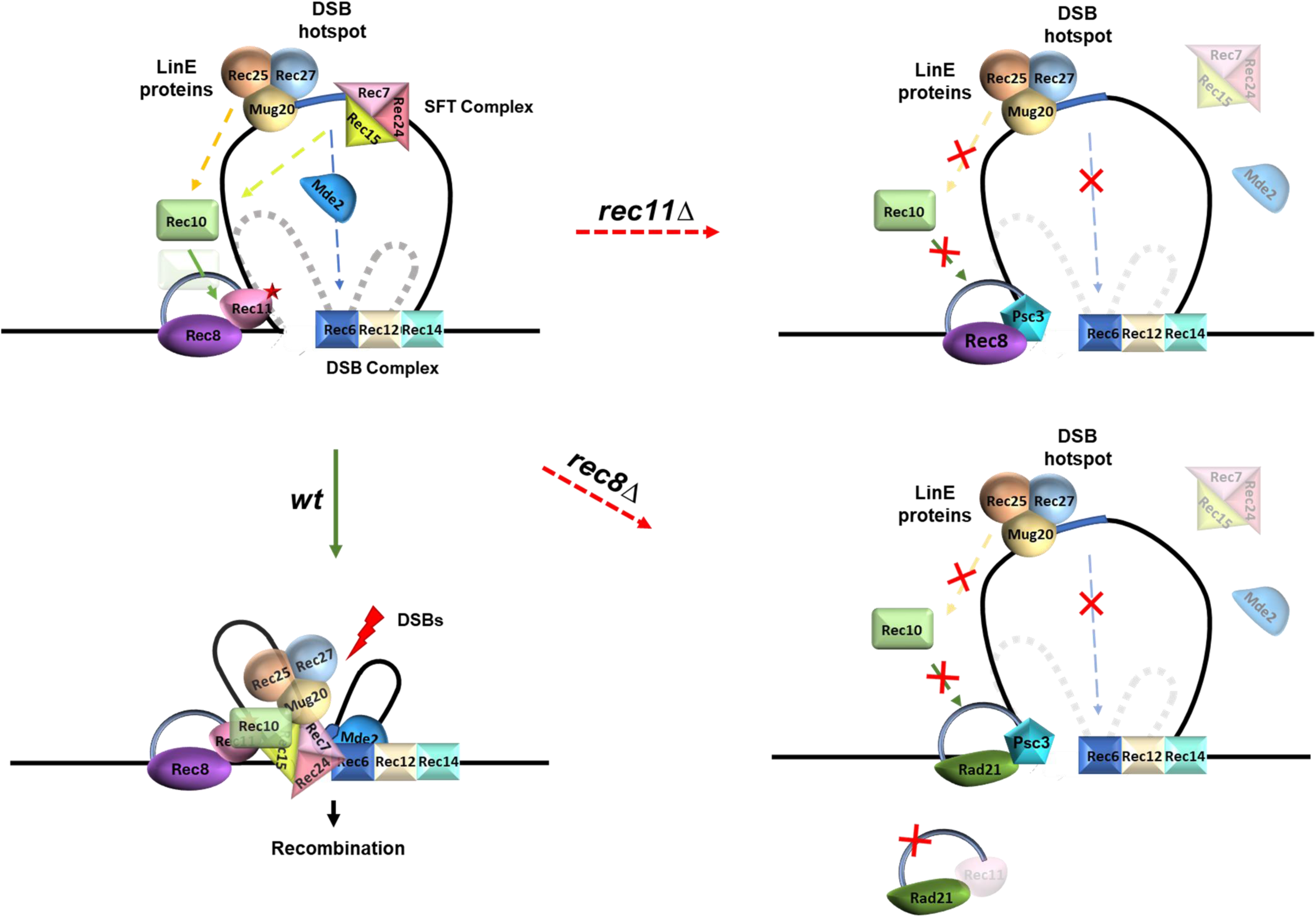
Model to explain the molecular basis for lack of activation of recombination in the absence of the meiotic cohesins. In the absence of Rec8, Rec11 levels are drastically downregulated resulting majorly in the formation of Rad21-Psc3 cohesin complex. Despite cohesin loading on the chromosome axes, there is lack of recruitment of the activator Rec10, which halts the activation of the DSB inducing Rec12 complex. In the absence of Rec11, Rec8-Psc3 complex takes over as the major cohesin complex and is enriched at the centromeres as well as the cohesin-rich axis. However, this complex also fails to recruit Rec10 to the linear elements/axes and impedes interaction with Rec15, thereby resulting in defective axial tethering of the hotspots in the loops.

In humans, RAD21L is a novel paralog of the kleisin subunit that is additionally present during meiosis and not seen in yeast ^28^. RAD21L and REC8 appear to occupy different positions on the axial elements of the synaptonemal complex, where RAD21L is thought to help bridge the non-sister chromatids near recombination intermediates ^29^. The two kleisins have distinct non-overlapping functions during meiosis and the exclusive expression of RAD21L1 only in prophase I till mid-pachytene suggests evolution of this paralog to perform only certain discrete functions that cannot be done by REC8. Perhaps RAD21L is an example of an end-point in molecular evolution of kleisins, wherein the redundant functions have largely been lost. Interestingly, in *REC8* or *RAD21L* double-knockouts with meiosis-specific SMC1β, mitotic RAD21 in complex with the mitotic SMC1α provides centromeric cohesion only during early prophase I, further highlighting the flexibility among paralogs^30^. In terms of meiotic functions, ectopic expression of SMC1α in SMC1β^-/-^, restores multiple defects associated with recombination and meiotic chromosome architecture, arguing for amount of cohesin being critical rather than the quality ^17^. In contrast, our work demonstrates that the composition of the complex is more important to determine the functional plasticity.

Another example in human somatic cells, are the paralogs STAG1 and STAG2 that are responsible for cohesion at the telomeres and centromeres, respectively and have distinct roles in DNA damage response, replication fork progression, gene regulation and genome organization ^31–36^. There is also synthetic lethality between the two paralogs due to their partial redundant functions that can be exploited for therapy in human cancers ^37,38^. All these studies highlight the extent of functional overlap between different cohesin paralogs. However, the mechanistic details of how these differences come about still remain unresolved.

### Why are the meiotic cohesin paralogs needed*?*

Reductional division in meiosis requires monopolar attachment of the sister kinetochores so that they move to the same pole at the end of meiosis I. This feature would need to be avoided during mitosis as it may result in uniparental disomy, as reported in several cancers that aberrantly express REC8 ^39^. Hence, the meiotic kleisin Rec8 can specifically get deposited at the centromere core and associate with factors such as MoaI to ensure mono-orientation of sister chromatids ^40,41^. Due to the two-step chromosome segregation, some basal level of cohesion is required during metaphase II, which led to the requirement of protecting centromeric cohesins from degradation at anaphase I. Interaction of Shugoshin-PP2A complex with Rec8 allows this protection, which seems to be missing in Rad21, thereby allowing equational division ^42,43^. Moreover, meiotic chromosomes are extensively reorganized leading to formation of axial elements and synaptonemal complex for homolog pairing stabilization, which would have necessitated the evolution of a modified subunit.

Distribution of recombination events such as crossovers and non-crossovers is tightly regulated and some chromosomal regions like the centromeres are actively repressed for recombination to avoid potential errors during segregation ^44–46^. Regulation of recombination can happen either at the DSB level or at the time of DNA repair. The distantly related yeast *S. cerevisiae,* has only one cohesin complex Rec8-Scc3^Psc3^ during meiosis, retaining the mitotic HAWK subunit. Unlike *S. pombe* where Rec8 is essential for induction of DSBs, *S. cerevisiae* doesn’t depend on cohesins to initiate Spo11- break formation but are critical for directing repair via homologous recombination ^47^. Although, centromeres in *S. cerevisiae* show presence of DSBs near centromeres, the enriched cohesins deposited in the flanking regions alter the type of DNA repair leading to absence of deleterious crossover events ^48^. On the contrary, *S. pombe* with larger centromeres containing heterochromatin, opted to suppress recombination by inhibiting induction of DSBs via altering the composition of the centromeric cohesins and biased the loading of Rec8-Psc3 complex that cannot activate recombination ^5,22^. Hence, allowing continued expression of the Psc3 in mitosis, unlike Rad21, provides flexibility to cells in utilizing the two complexes differentially to regulate spatial distribution of recombination events. Moreover, reducing Rec11 expression when Rec8 is absent, results in lowered chance of Rad21-Rec11 complex to be available for loading in the chromosomal arms to activate DSBs (Fig. 6). Overall, our work highlights how systematic determination of overlapping and distinct functions of paralogs in mitosis and meiosis, may provide insights regarding the order of events during evolution of these processes in eukaryogenesis.

## Materials and methods

### Strains and allele generation

The *S. pombe* genotypes generated and used in this study are described in Table S1. Strains were generated by crossing appropriate *h+* and *h-* strains followed by random spore analysis^49^. All *pat1-114* strains were grown at 25°C, while the rest were cultured at 30°C. Oligomers used in the study are described in Table S2.

#### rad21-rec11 fusion allele

*FLAG-ura4+* cassette was replaced in the allele *rec8D::Padh1-rad21+-FLAG-ura4+* and *rec8D::Prec8-rad21+-FLAG-ura4+* (procured from the NBRP) with *rec11-HA-KanR* cassette via homologous recombination after yeast transformation. A 27 aa flexible linker (KESGSVSSEQLAQFRSLDEGKSSGSGS) was added between *rad21* and *rec11* ^50^. Overlap PCR was done to construct *rad21(3’end)-linker-rec11-HA-KanR-rec8(3’UTR)*. MO440 and MO442 were first used to amplify *rec11-HA-KanR* ({rom gDNA containing *ura4+-Padh1-rec11-3HA-kanMX6* (MN lab)} with 80 bp homology from *rec8*-3’UTR while adding a 39 bp linker to *rec11*-5’. The purified PCR product was then amplified using MO439 and MO442 to add another 42 bp to the linker. Finally, MO441 and MO442 were used to amplify the 2nd PCR product adding 80 bp homology from *rad21*. The purified amplicon was then used for replacing the *FLAG-ura4+* cassette. KanR transformants were selected and the integration was confirmed by absence of growth in minimal media without uracil.

#### rad21 overexpression (oe) allele

*ura4+-Padh1* cassette was amplified from FYP4963 (procured from NBRP) using MO217 and MO218 while adding a 20 bp linker upstream of *ura4+* and 18 bp linker downstream of *Padh1*. 400 bp *rad21* homology regions were amplified using the primer pairs MO215, MO216 and MO219, MO136. The homology regions were then added to the *ura4+-Padh1* cassette on either side via overlap PCR. The fragment was then transformed to replace the native promoter of *rad21*(∼1100 bp before Rad21 start codon). The integration was confirmed by colony PCR using MO90 and MO136.

#### rec15-myc and mug20-myc alleles

*3xMyc-linker-hygR* allele having 18 bp linker attached upstream of 3xMyc and 21 bp linker attached downstream of *hygR* was amplified from gDNA of strain containing *swi6-myc-HygR* (MN lab). MO415, MO417 for *rec15* and MO419, MO421 for *mug20* were used to amplify the allele while adding 41bp homology to the left and 38bp homology to the right of the cassette. In total ∼80 bp homology regions corresponding to *rec15* and *mug20* were added via overlap PCR using the primer pairs MO416, MO418 and MO420, MO422, respectively. The amplicon was then inserted in the 3’-UTR of *rec15* and *mug20* via homologous recombination after yeast transformation. The integration was confirmed using MO113 and MO430 for *rec15* and MO8 and MO100 for *mug20*.

#### rec11-gbp (gfp binding protein) allele

*rec11* was amplified using MO493 and MO494 containing restriction enzyme sites. The fragment was cloned between KpnI and BamHI in FYP3976 (procured from NBRP). The confirmed plasmid was then digested with ApaI and then transformed into strains containing *rad21-gfp*. The transformants were selected based on Hygromycin resistance and further confirmed by absence of growth in minimal media lacking lysine ^51^.

### *S. pombe* transformation

Lithium acetate transformation method was used as described previously ^52^. ∼1X10^7 log phase cells were harvested and resuspended in TE/LiOAc buffer after washing and incubated at 25°C for 30 min. The cells were then pelleted and resuspended in TE/LiOAc with 40% PEG 4000, 50ug salmon sperm DNA (Invitrogen 15632011), and 1µg PCR product. The cells were then incubated at 25°C for 30 min. DMSO was added and the cells were incubated at 42 °C for 20 min. The cells were then collected and allowed to recover in appropriate media for 18-22 h. After plating on selection media, the transformants were isolated and the colonies were screened via PCR for confirmation.

### Meiotic induction and FACS

The strains were first cultured in YEL till saturation and then subcultured in supplemented EMM2 +NH4Cl. This subculture was used to set up a secondary culture of larger volume. Once the culture reached 0.3-0.4 OD_600_, the cells were shifted to EMM2 -NH4Cl to starve the cells for 15 h. 2.5% v/v 20% NH_4_Cl was added to the cultures and the temperature was shifted to 34°C to induce meiosis in *pat1-114* strains. Meiotic induction was then confirmed via FACS. For checking the expression of proteins, the cells were harvested at 0, 3, 3.5, 4 and 6 h post meiotic induction. For ChIP qPCR experiments, the samples were collected at 3.5 h post-meiotic induction.

### Immunoblotting

Post-meiotic induction, the cells were harvested at 0, 3, 3.5, 4 and 6 h time points. The cells were then washed and lysed using 1.85M NaOH in a reducing environment. 50% Trichloroacetic acid (TCA) was used to precipitate out the proteins. The pellet was then resuspended in 1X Laemmli buffer, followed by neutralisation with pH-8 Tris-HCl. The protein lysate was loaded on 7.5% SDS-PAGE gel after equating and then transferred onto PVDF membrane. A 5% skimmed milk solution was used for blocking the membrane followed by probing using antibodies against the respective tags {anti-HA (Invitrogen 26183), anti-FLAG (Sigma F1804), anti-V5 (Invitrogen R96025), anti-Myc (Invitrogen MA121316), anti-Gapdh (Invitrogen MA5-15738) and anti-Rad21 (BioAcademia Inc 63-139)}. Goat anti-mouse IgG H+L (Jackson Immunoresearch, 115-035-003) was used as the secondary antibody and the blots were developed using Chemiluminescent substrate (Thermo Fisher Scientific, 34580). The blots were imaged using iBright CL1500 imaging system.

### Chromatin Immunoprecipitation

Cells were harvested at 3.5 h post meiotic induction. The cells were fixed using 1.1% crosslinking solution (1.1% formaldehyde (Invitrogen 28908), 5mM HEPES-KOH, 550mM NaCl and 0.1mM EDTA), the excess formaldehyde was quenched by 250mM glycine solution. The cells were washed twice in ice-cold TBS. The pellet was then lysed with the help of liquid nitrogen upon resuspending in 500µl of Breaking buffer (100mM Tris-Cl pH-8, 20% glycerol and 1mM PMSF). The lysate was collected in microfuge tube and washed with FA buffer (50mM HEPES-KOH, 150mM NaCl, 1mM EDTA, 1% Triton X-100, 0.1% Sodium deoxycholate and 1X Protease inhibitor cocktail). 300µl FA buffer was used to resuspend the pellet followed by sonication Diagenode Biruptor for 25 cycles (30sec on/off at high amplitude). The volume was made up to 1.5 ml with FA buffer and the supernatant was collected upon centrifugation. A small volume of lysate was kept aside to be processed as input. The remaining lysate was precleared by incubating with magnetic beads (Invitrogen 1003D) for 4 h at 4 °C. Precleared lysate pertaining to ∼3 µg of DNA was incubated with the respective antibody overnight at 4 °C. The antibody bound Protein-DNA was then incubated with magnetic beads for 4 h at 4 °C. The beads were then washed twice each with FA buffer (50mM HEPES-KOH, 150mM NaCl, 1mM EDTA, 1% Triton X-100, 0.1% Sodium deoxycholate), FA-HS buffer (50mM HEPES-KOH, 500mM NaCl, 1mM EDTA, 1% Triton X-100, 0.1% Sodium deoxycholate), RIPA buffer (10mM Tris pH-8, 0.25M LiCl, 0.5% NP40, 0.5% Sodium deoxycholate and 1mM EDTA) and TE buffer (10mM Tris pH-8 and 1mM EDTA). 2X STOP buffer (20mM Tris pH-8, 100mM NaCl, 20mM EDTA and 1%SDS) was used for elution twice at 65 °C for 30 min. The elute and 10% input were decrosslinked by incubation overnight at 65 °C followed by RNAse and Proteinase treatment at 37 °C, 2 h each. The DNA was then purified using a PCR purification kit (Qiagen 28115). TB Green Premix Ex Taq II buffer mix (Takara RR820A) was used to set up the qPCR. The data was analysed using GraphPad Prism version 10.6.1 (892). The oligomers used for ChIP qPCR are mentioned in the Table S2.

### Fluorescence-based tetrad assay for chromosomal segregation

The parental strains were spotted on SPA media for 2-3 days at 25°C and meiosis was confirmed via asci formation. The mating mix was then put on a slide with 20 µl 50% glycerol. Leica DM6 upright epifluorescence microscope was used for imaging the asci, Texas Red (TxR) filter to detect Tdtomato and SBL filter to detect mCerulean using a 63X objective. Atleast 200 asci were imaged per strain. Fiji and LASX software were used for processing and analysing the images. The frequencies for each class of segregation profiles were calculated and plotted. A minimum of 3 biological replicates were performed for each genotype, unless otherwise mentioned.

### Genetic recombination assay

Recombination across the arms was measured across the *ade6-arg1* interval using prototrophic markers. Recombination across *cen3* was measured using prototrophic/antibiotic markers (*hygR* and *his3+)* introduced at *chk1* and near *mid1,* respectively. Recombination frequency was determined by random spore analysis. Recombination frequency was calculated as the percentage of recombinant colonies over the total number of colonies screened and the mean was plotted as bar graphs. For each genotype, the data were collected for at least 3 independent crosses (biological repeats). Error bars for all data represent SEM. The data for individual experiments are provided in Tables S5 and S6.

### AlphaFold analysis

Sequences pertaining to specific domains of cohesins were fed into AlphaFold 3 based on the annotations in Pombase. The 0th model obtained were then aligned using Chimera ^19,53,54^.

### Statistical Analysis

Two- tailed Fisher’s exact test was used to analyse the data for genetic recombination assays. Two-way ANOVA test (Tukey’s multiple comparisons test) was used to analyse the fluorescence-based tetrad assay and ChIP qPCR data. All the analysis was done using GraphPad Prism version 10.6.1 (892).

## Acknowledgement

We thank Nikhil Venketesh, Laxmi Sharma, Reva Deokule, Rupal Yadav and Shubhangi Sharma for technical help. We thank Dr. Girish Deshpande and MN lab members for their inputs on the manuscript and helpful discussions throughout the course of this study. We acknowledge Prof. Gerald R. Smith for sharing *S. pombe* strains and Prof. Alexander Lorenz for sharing the spore-autonomous mCerulean and tdTomato expressing *S. pombe* strains UoA726 and UoA727. Purva acknowledges Prime Minister’s Research Fellowship (0702610). This study was supported by the Science and Engineering Research Board (SERB) - Promoting Opportunities for Women in Exploratory Research (POWER) Research grant [SPG/2022/000881] and intramural funds to MN. We also thank IISER Pune shared facilities including those for Microscopy and FACS for infrastructural support.

## Author Contributions

Purva and M.N. conceived the study, designed the experiments and analysed the data. Purva and A.D. performed the experiments. M.N. acquired resources and funding and wrote the original draft of the manuscript with inputs from other authors.

