## Supplementary Data for "Molecular mechanism for the functional divergence between cohesin paralogs during meiosis"

**A**

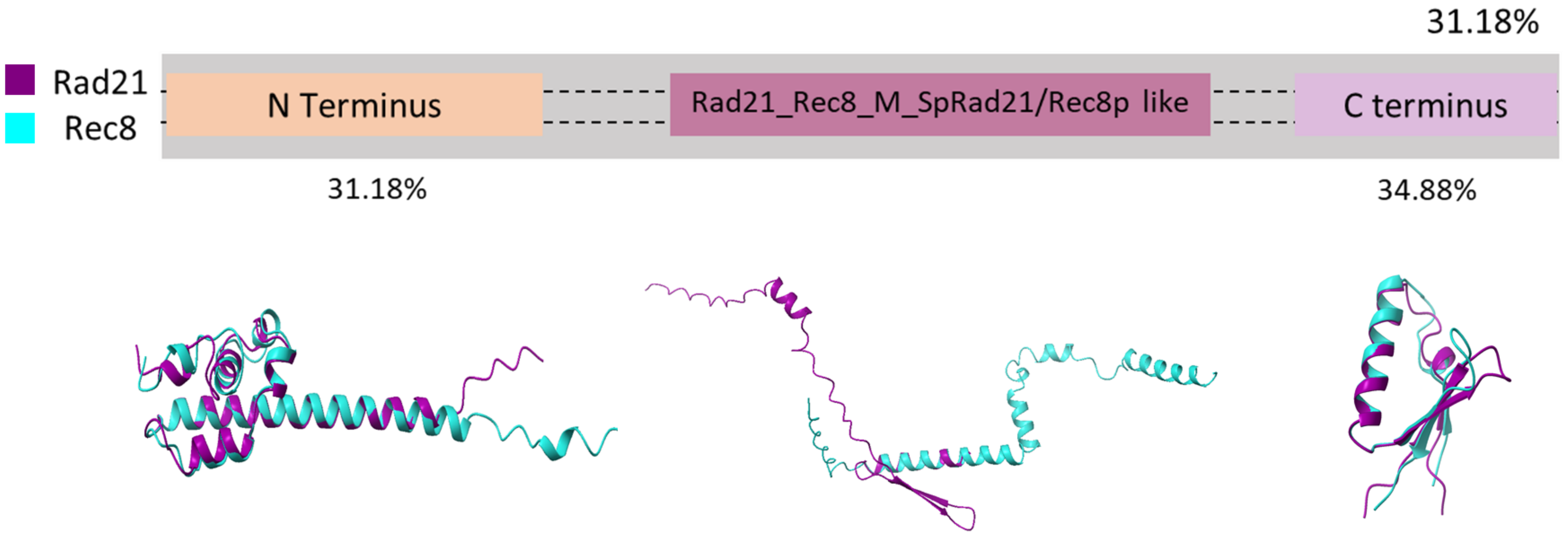


| | | | | |
|---|---|---|---|---|
| Rad21 | 0.78 | 0.3 | 0.7 | pTM score |
| Rec8 | 0.73 | 0.27 | 0.6 | |

**B**

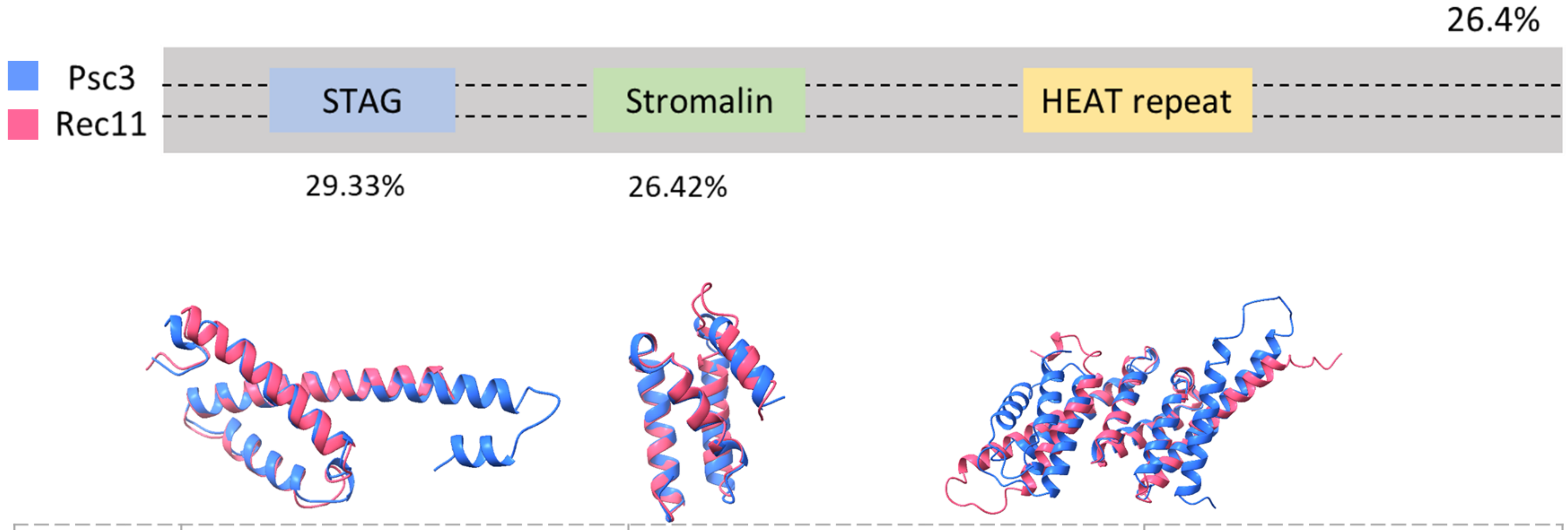


| | | | | |
|---|---|---|---|---|
| Psc3 | 0.81 | 0.86 | 0.84 | pTM score |
| Rec11 | 0.72 | 0.73 | 0.71 | |

Figure S1

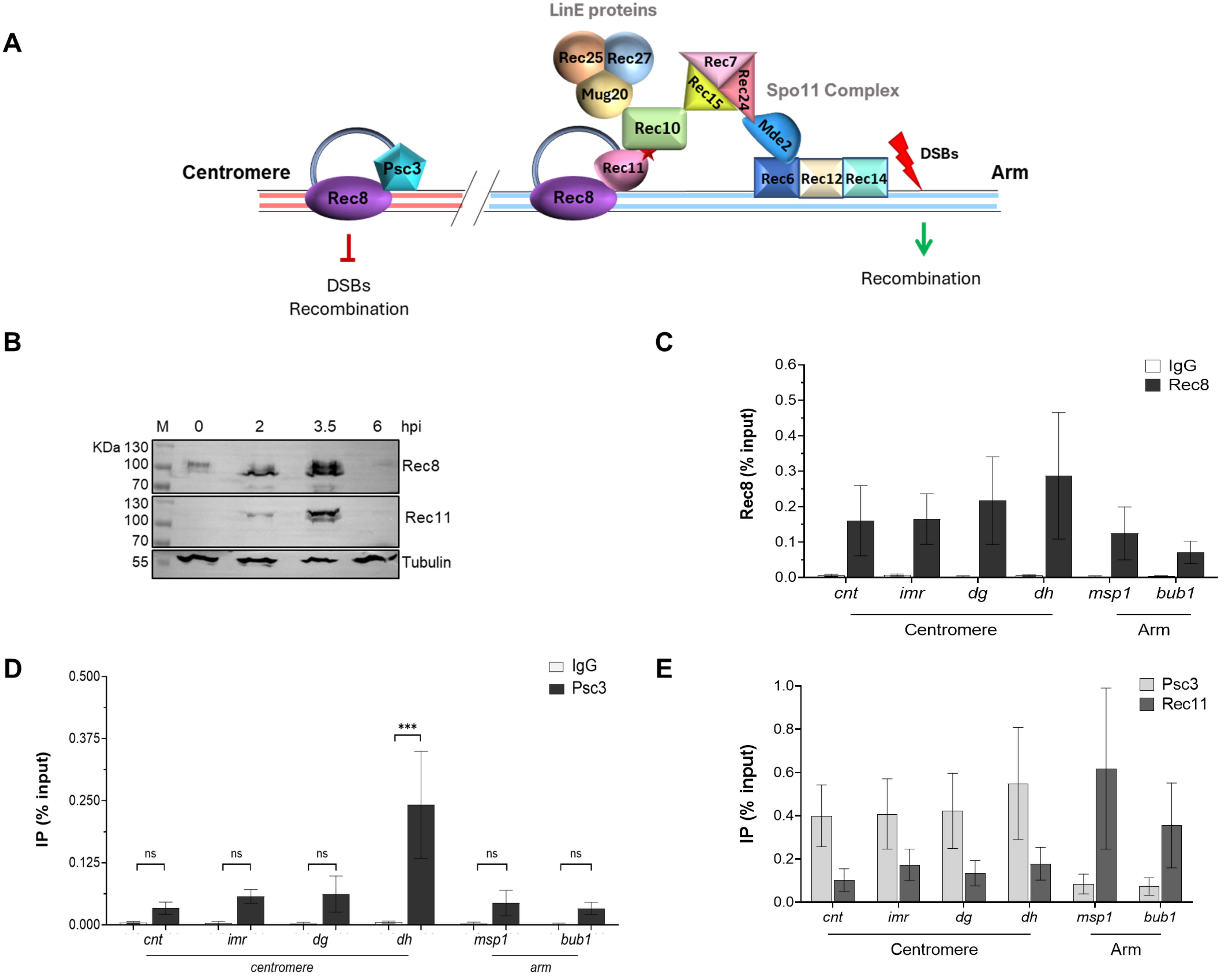
A
LinE proteins
Rec25
Rec27
Mug20
Rec10
Rec7
Rec15
Rec24
Spo11 Complex
Mde2
Psc3
Rec11
Rec6
Rec12
Rec14
DSBs
Centromere
Rec8
Rec8
Arm
DSBs
Recombination
Recombination
B
M
0
2
3.5
6
hpi
KDa 130
100
70
Rec8
130
100
70
Rec11
55
Tubulin
C
Rec8 (% input)
IgG
Rec8
cnt
imr
dg
dh
msp1
bub1
Centromere
Arm
D
IP (% input)
IgG
Psc3
ns
***
cnt
imr
dg
dh
msp1
bub1
centromere
arm
E
IP (% input)
Psc3
Rec11
cnt
imr
dg
dh
msp1
bub1
Centromere
Arm

Figure S2

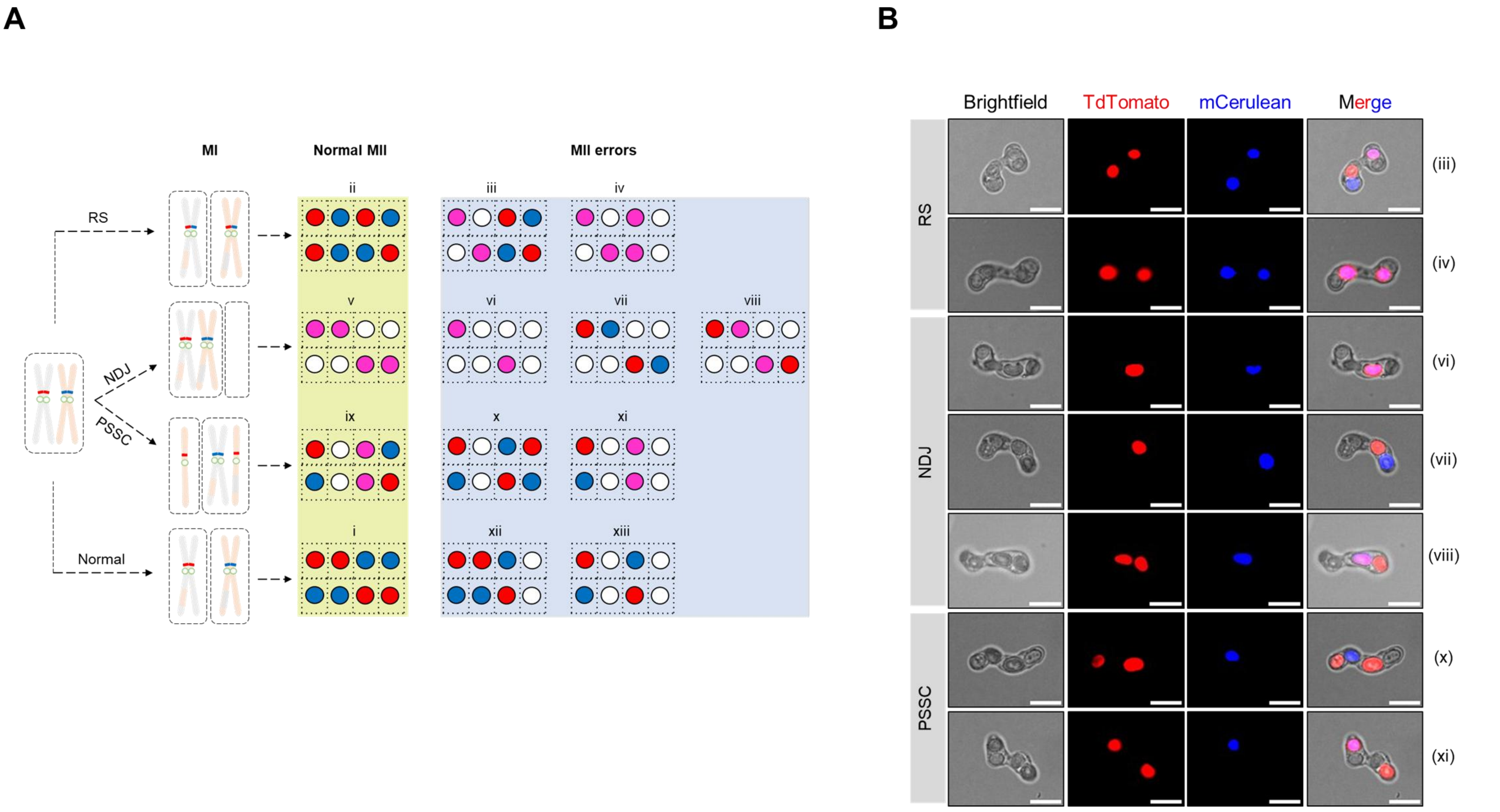
A
MI
Normal MII
MII errors
RS
NDJ
PSSC
Normal
ii
iii
iv
v
vi
vii
viii
ix
x
xi
i
xii
xiii
B
Brightfield
TdTomato
mCerulean
Merge
RS
NDJ
PSSC
(iii)
(iv)
(vi)
(vii)
(viii)
(x)
(xi)

Figure S3

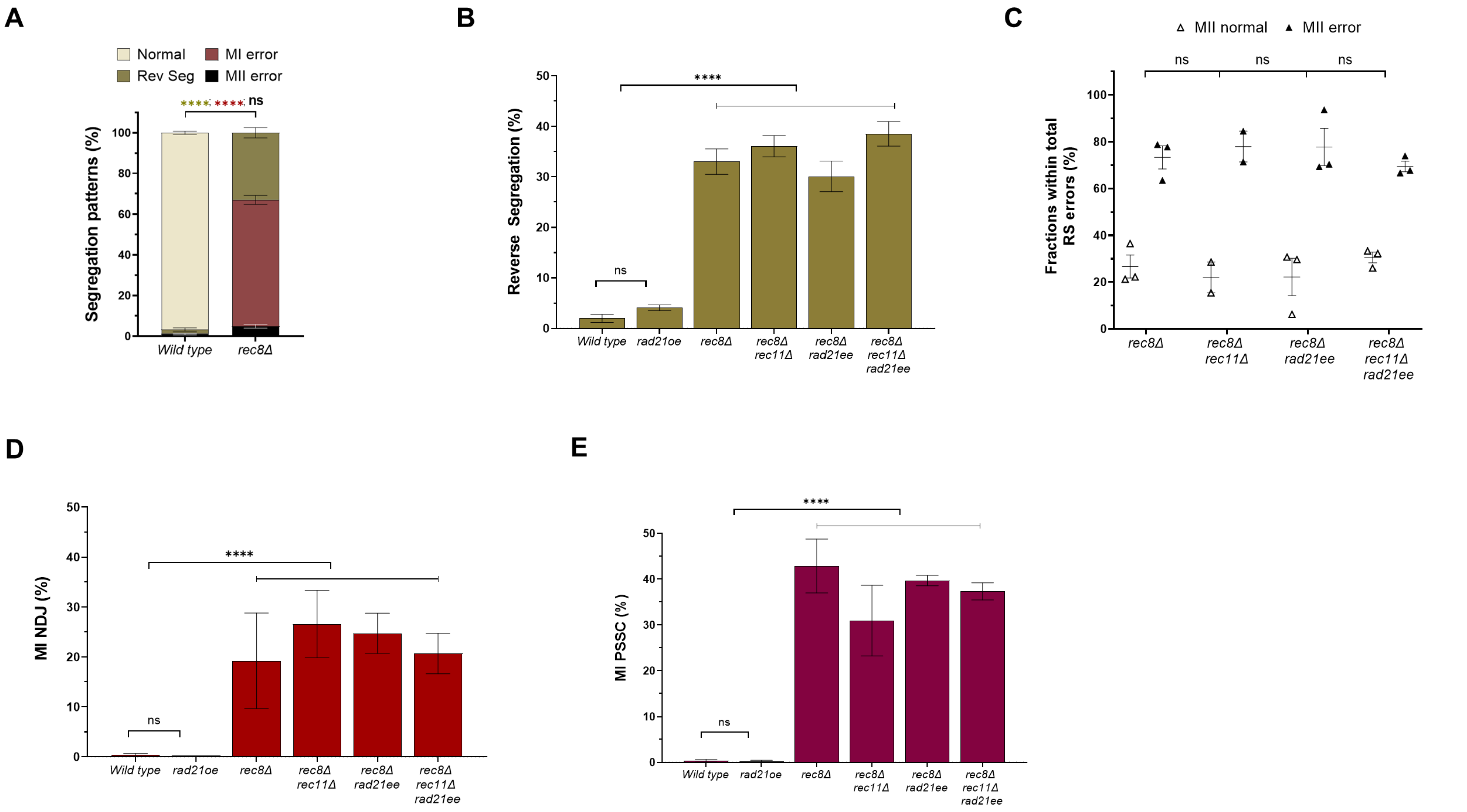
A
Normal
MI error
Rev Seg
MII error
****
****
ns
Segregation patterns (%)
Wild type
rec8Δ
B
****
ns
Reverse Segregation (%)
Wild type
rad21oe
rec8Δ
rec8Δ rec11Δ
rec8Δ rad21ee
rec8Δ rec11Δ rad21ee
C
MII normal
MII error
ns
ns
ns
Fractions within total RS errors (%)
rec8Δ
rec8Δ rec11Δ
rec8Δ rad21ee
rec8Δ rec11Δ rad21ee
D
****
ns
MI NDJ (%)
Wild type
rad21oe
rec8Δ
rec8Δ rec11Δ
rec8Δ rad21ee
rec8Δ rec11Δ rad21ee
E
****
ns
MI PSSC (%)
Wild type
rad21oe
rec8Δ
rec8Δ rec11Δ
rec8Δ rad21ee
rec8Δ rec11Δ rad21ee

Figure S4

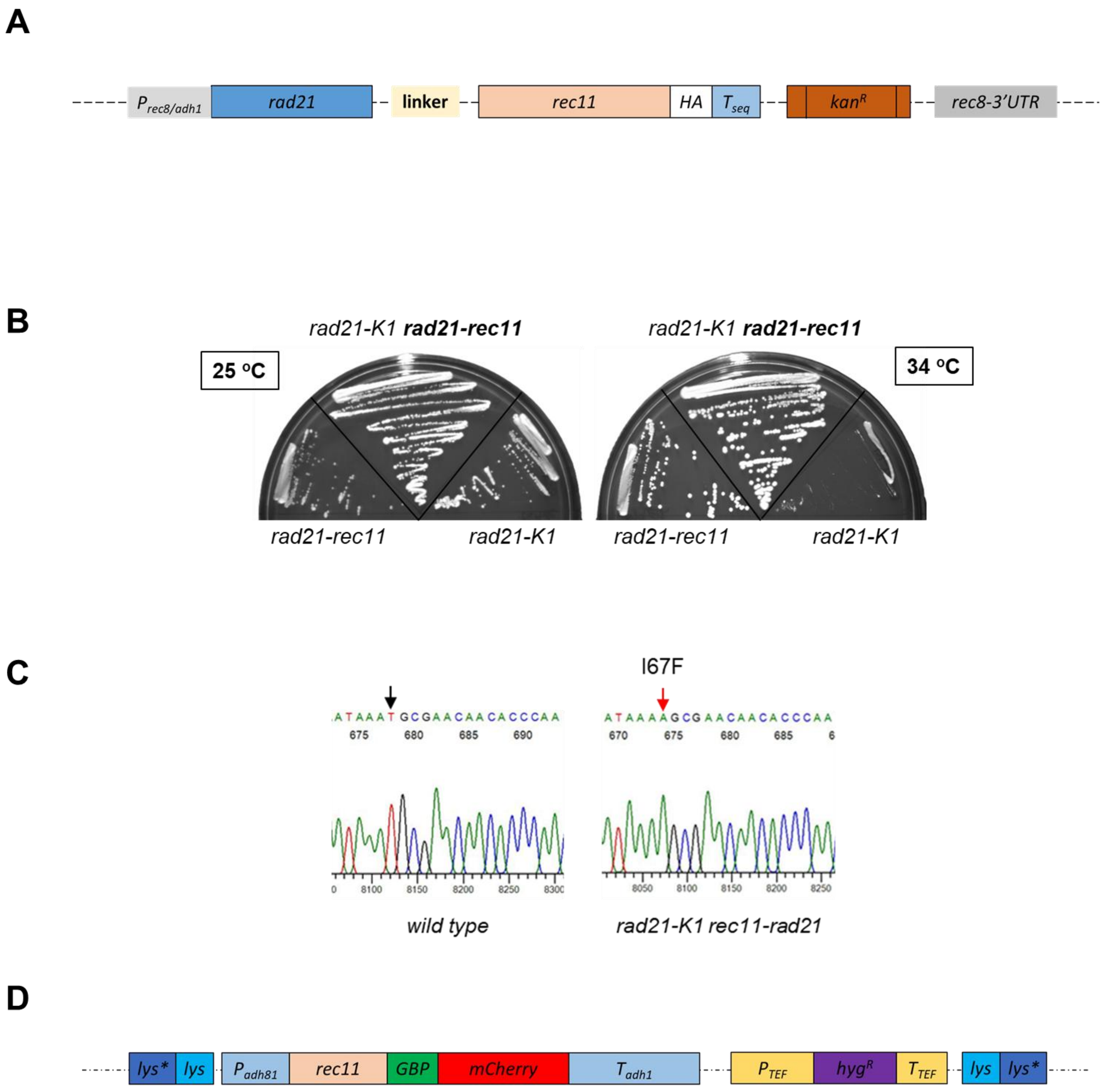
A
P rec8/adh1
rad21
linker
rec11
HA
T seq
kanR
rec8-3'UTR
B
rad21-K1 rad21-rec11
rad21-K1 rad21-rec11
25 °C
34 °C
rad21-rec11
rad21-K1
rad21-rec11
rad21-K1
C
I67F
ATAAATGCGAACAACACCCAA
675 680 685 690
ATAAAAGCGAACAACACCCAA
670 675 680 685
8100 8150 8200 8250
8050 8100 8150 8200 8250
wild type
rad21-K1 rec11-rad21
D
lys*
lys
P adh81
rec11
GBP
mCherry
T adh1
P TEF
hygR
T TEF
lys
lys*

Figure S5

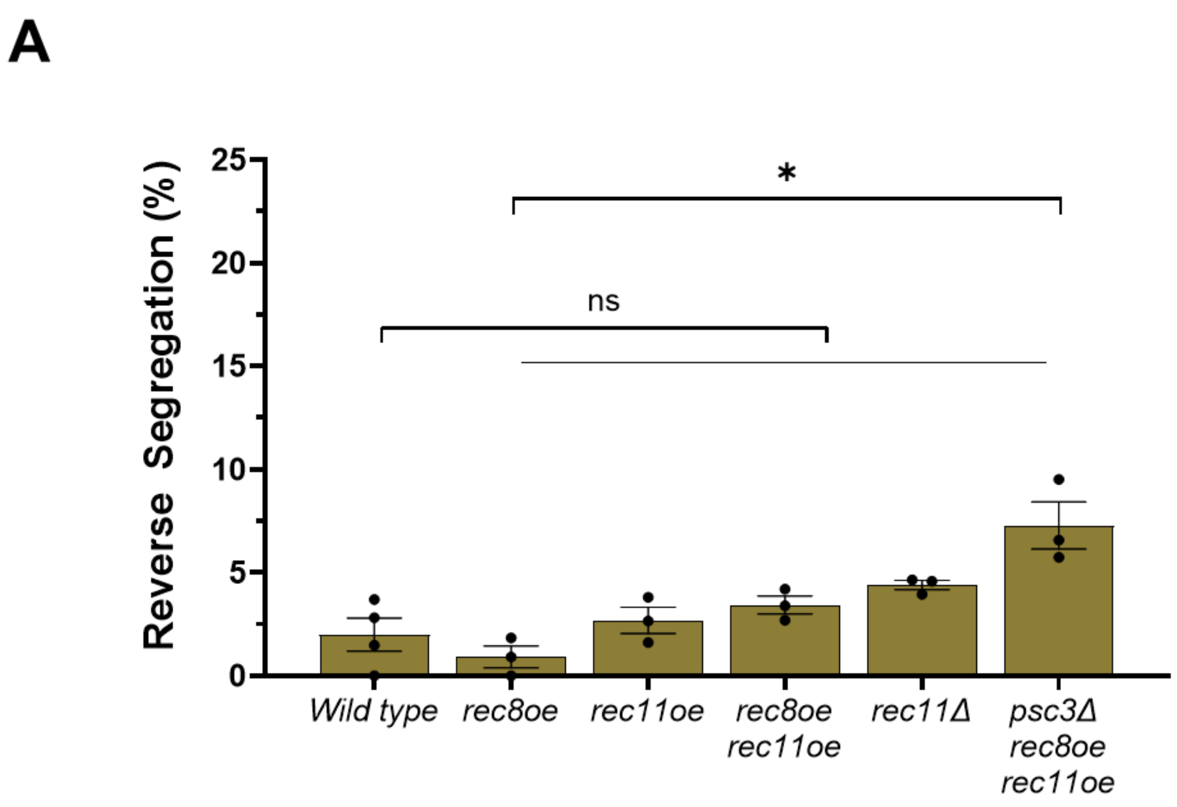
A
Reverse Segregation (%)
25
20
15
10
5
0
*
ns
Wild type
rec8oe
rec11oe
rec8oe rec11oe
rec11Δ
psc3Δ rec8oe rec11oe

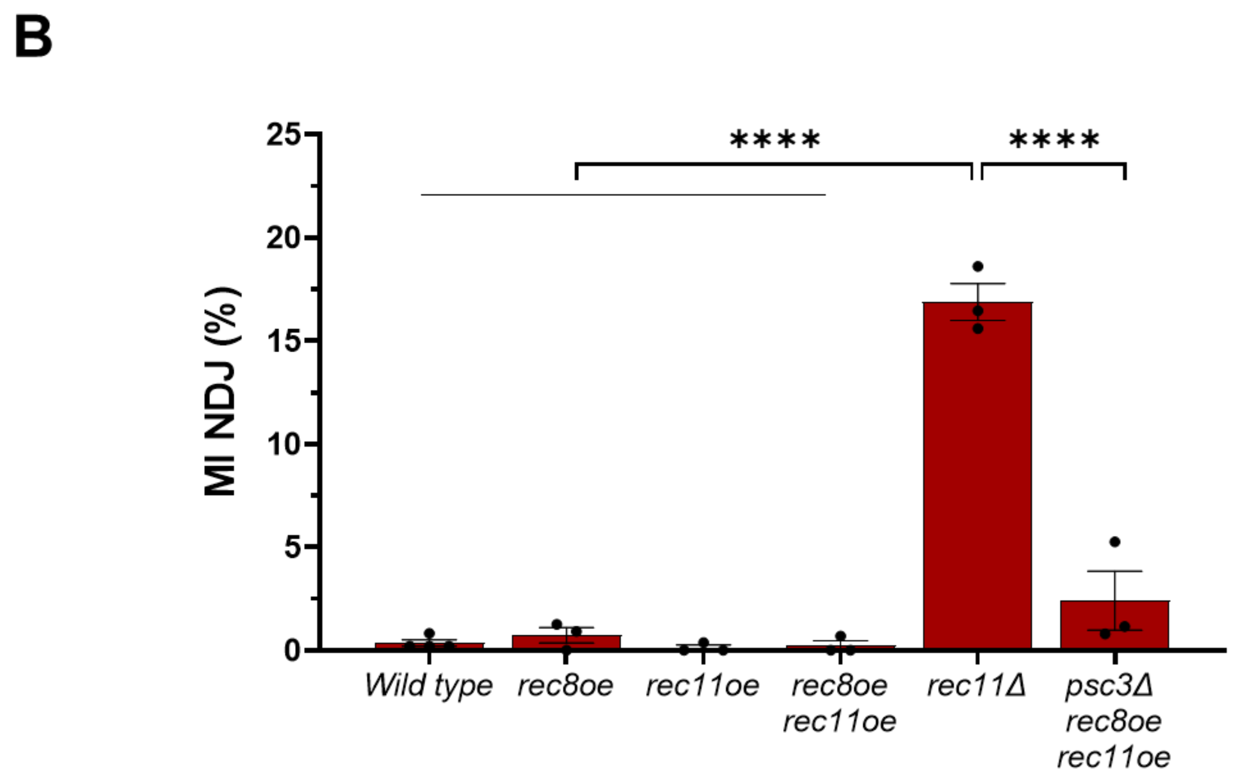
B
MI NDJ (%)
25
20
15
10
5
0
****
****
Wild type
rec8oe
rec11oe
rec8oe rec11oe
rec11Δ
psc3Δ rec8oe rec11oe

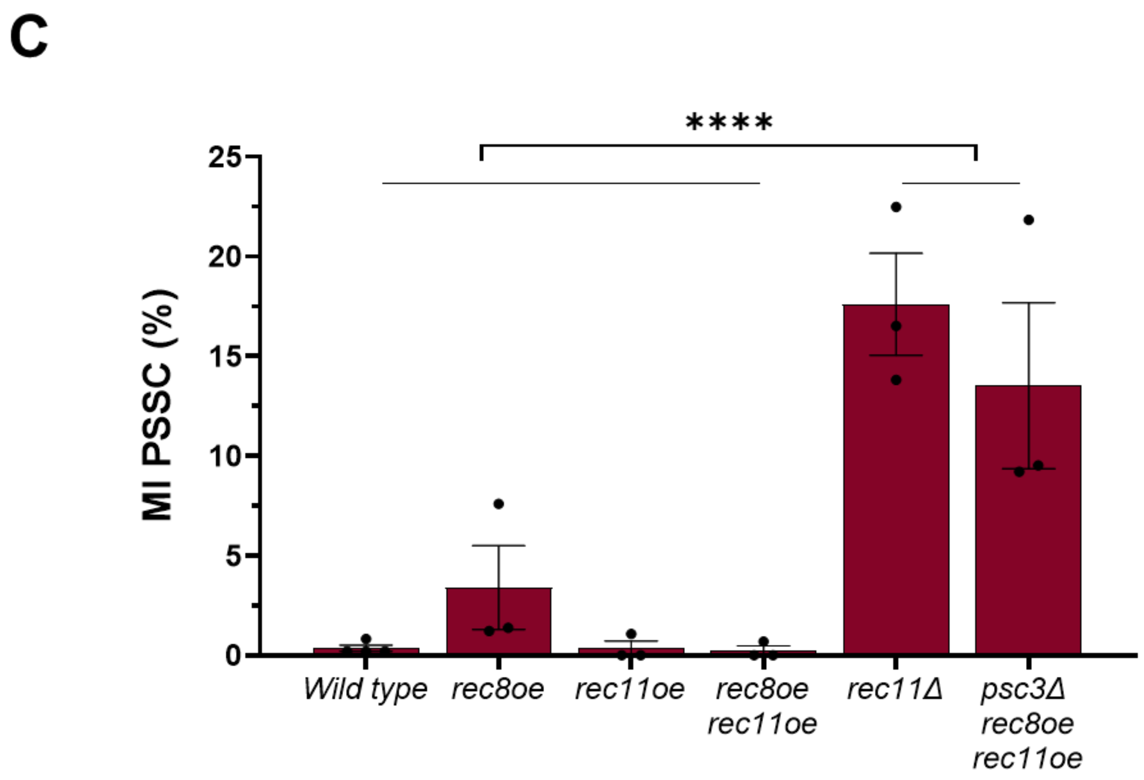
C
MI PSSC (%)
25
20
15
10
5
0
****
Wild type
rec8oe
rec11oe
rec8oe rec11oe
rec11Δ
psc3Δ rec8oe rec11oe

Figure S6

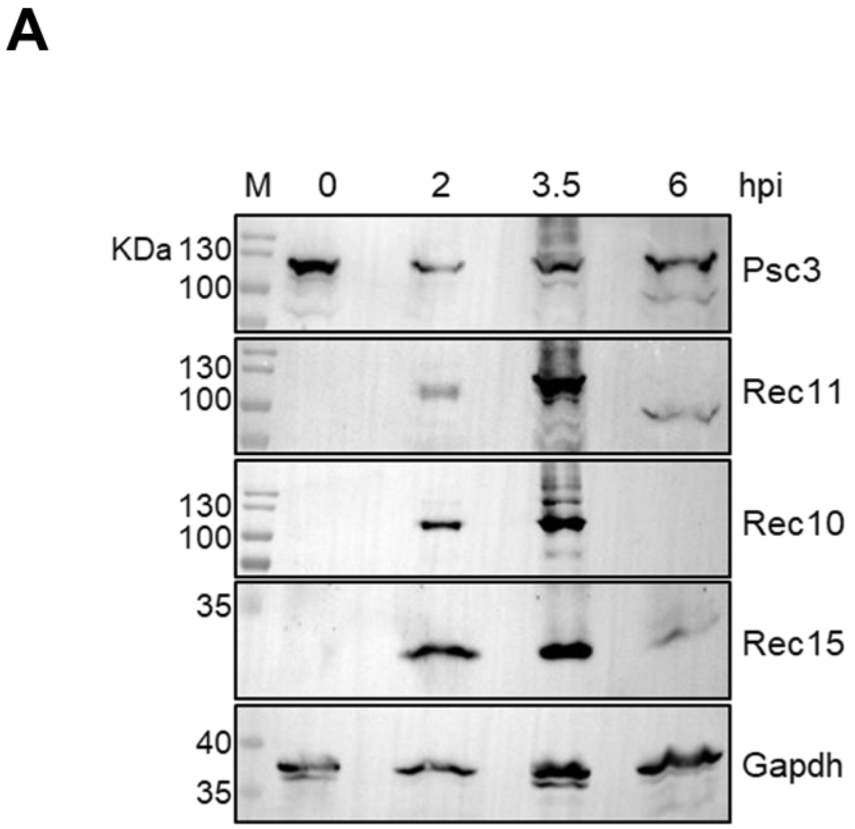
A
M
0
2
3.5
6
hpi
KDa 130
100
Psc3
130
100
Rec11
130
100
Rec10
35
Rec15
40
35
Gapdh

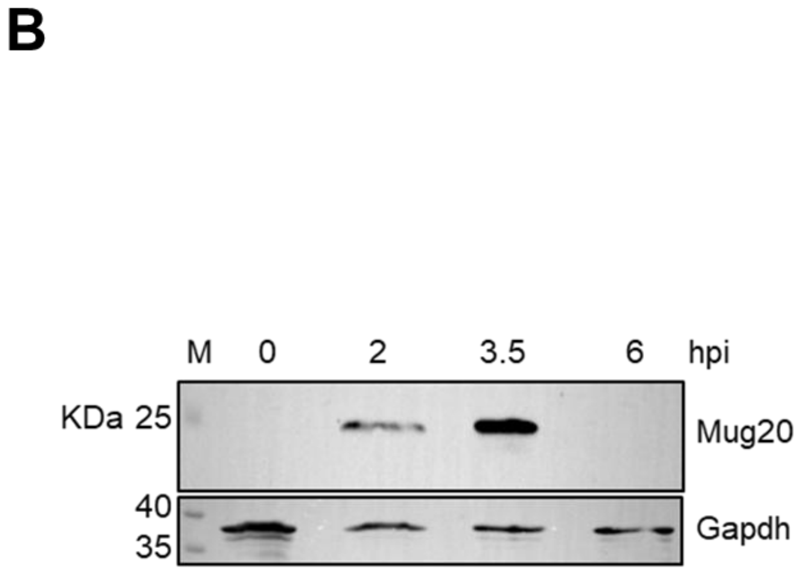
B
M
0
2
3.5
6
hpi
KDa 25
Mug20
40
35
Gapdh

Figure S7

**Supplementary Figure Legends**

**Figure S1. Sequence similarity and Alpha Fold predicted structural similarity between the Kleisin and HAWK cohesin subunits. A.** Percentage of the amino acid sequence homology between Rad21 and Rec8 at the conserved N-terminus, C-terminus and a central domain that interacts with the HAWK partner calculated using sequence information from PomBase via BLAST. **B.** Percentage of the amino acid sequence homology between Psc3 and Rec11 at the conserved STAG domain, Stromalin domain and the HEAT repeat associated domain, which interacts with the Kleisin partner, calculated using BLAST. The overlapped structures are shown for all the domains. pTM scores for the AlphaFold predictions are mentioned.

**Figure S2. Distribution of the cohesin subunits during meiosis across centromere and chromosomal arms. A.** Schematic diagram showing the type of cohesins occupying the centromeric and arm regions during meiosis. This dictates whether DSBs are induced at the given region followed by activation of homologous recombination to repair the breaks. **B.** Expression profile via western blotting for the meiotic cohesins Rec8 and Rec11 in *pat1-114* induced meiosis at various time points post-induction. **C-E.** ChIP qPCR for Rec8 (C), Psc3 (E) and Rec11 (E) in wild type at centromere (*cnt, imr, dg* and *dh*) and arm (*msp1* and *bub1*) loci. ChIP qPCR for Psc3 (D) in wild type and *rec8*Δ at the same loci. Pulldowns with IgG were used as controls. The bars represent the mean of at least three independent experiments performed in triplicates; error bars=SEM. ***p <0.001; ns p >0.05. The data were analyzed using Two-way ANOVA test (Tukey's multiple comparisons test).

**Figure S3. Tetrad assay for segregation fidelity to classify diverse types of meiotic errors. A.** Schematic representation of the fluorescence-based tetrad assay employed for determining segregation fidelity and the various types of patterns that can be obtained corresponding to the errors in specific stages of meiotic division. Consequences of normal vs erroneous MII for each case is also shown. Patterns arising from mirroring each or both halves of the asci are not shown. **B.** Representative images for reverse segregation (RS), meiosis I non-disjunction (MI NDJ) and meiosis I premature separation of sister chromatids (MI PSSC) error types that were not included in Fig. 2A are shown along with the corresponding numbers as depicted in the schematic cartoon in panel A. Scale bars represent 10 μm.

**Figure S4. Distribution of the subtypes of segregation errors in MI for *rec8*Δ and similar genotypes. A.** Plot showing frequency of normal, RS, MI and MII segregation errors in wild type and *rec8Δ* genotypes. **B, D-E.** Bar graph showing total percentage of RS (B), MI NDJ (D) and MI PSSC (E) error events among all the segregation error types in wild type*, rad21oe, rec8Δ, rec8Δ rec11Δ, rec8Δ rad21ee* and *rec8Δ rec11Δ rad21ee* genotypes. **C.** Graph showing fraction of normal and erroneous MII segregation within RS events in *rec8Δ, rec8Δ rec11Δ, rec8Δ rad21ee* and *rec8Δ rec11Δ rad21ee* genotypes. All data are mean ± SEM (n $\geq$ 3 experiments; 2 for *rec8Δ rec11Δ*), assaying 563, 506, 362, 208, 438 and 395 tetrads for wild type*, rad21oe, rec8Δ, rec8Δ rec11Δ, rec8Δ rad21ee* and *rec8Δ rec11Δ rad21ee*, respectively. ****p <0.0001; ns p >0.05. The data were analyzed using Two-way ANOVA test (Tukey's multiple comparisons test). The data for individual experiments are provided in Table S3.

**Figure S5. Strategies to force Rad21 and Rec11 in a complex during meiosis. A.** Schematic showing generation of the *rad21-rec11oe* fusion allele. **B.** Growth of *rad21-rec11oe, rad21-k1* and *rad21-k1 rad21-rec11oe* genotypes at permissive (25 $^{o}$C) and restrictive (34 $^{o}$C) temperatures. **C.** Sequence confirmation for the point mutation in the *rad21-k1* allele in the *rad21-k1 rad21-rec11oe* strain that leads to the I67F change responsible for the temperature sensitivity of the *rad21-k1* allele. **D.** Schematic showing the allele for *rec11-gbp* (GFP binding protein) that would allow it to bind to Rad21-GFP in cells to force complex formation.

**Figure S6. Distribution of the subtypes of segregation errors in MI for *rec11*∆ and *psc3*∆ genotypes. A-C.** Bar graph showing total percentage of RS (A), MI NDJ (B) and MI PSSC (C) error events among all the segregation error types in wild type*, rec8oe, rec11oe, rec11oe rec8oe, rec11Δ* and *psc3Δ rec8oe rec11oe* genotypes. All data are mean ± SEM (n $\geq$ 3 experiments), assaying 563, 459, 659, 460, 614 and 289 tetrads for wild type*, rec8oe, rec11oe, rec11oe rec8oe, rec11Δ* and *psc3Δ rec8oe rec11oe,* respectively. ****p <0.0001; *p<0.05; ns p >0.05. The data were analyzed using Two-way ANOVA test (Tukey's multiple comparisons test). The data for individual experiments for panels are provided in Table S4.

**Figure S7. Meiotic expression profiles for the proteins involved in the meiotic recombination pathway. A-B.** Expression profile via western blotting for the HAWK cohesin subunits Psc3 (A), Rec11 (A) recombination proteins Rec10 (A), Rec15 (A) and DSB hotspot determinant Mug20 (B) in *pat1-114* induced meiosis at various time points post-induction. The blots were probed with anti-FLAG (for Psc3), anti-HA (for Rec11), anti-V5(pk) (for Rec10), anti-Myc (for Rec15 and Mug20) and anti-Gapdh antibodies. These tagged proteins were used for the ChIP-qPCR analysis for determining the distribution of these proteins across chromosomal loci during meiosis in wild type and *rec11*∆ genotypes.

**Table S1 - List of *Schizosaccharomyces pombe* strains used in the study.**

| Strain # | Genotype | Source | Used in Figure # |
|---|---|---|---|
| MP4 | *ade6-M210* | GRS lab | 2E, 3A,4A |
| MP41 | *ade6-52 ura4-D18 rec11::kanR his3-D1* | GRS lab | 4A |
| MP96 | *ade6-52 arg1-14* | GRS lab | 2E, 3A, 4A |
| MP172 | *ade6-52 ura4-D18 his3-D1 chk1::hygR mid1::his3+* | GRS lab | 2E, 4A |
| MP208 | *CEN1::his3+-PSPOG_00147-tdTomato his3-D1 ura4-D18* | Lorenz lab | 2B-D, 4B, S4A |
| MP209 | *ura4-D18 his3-D1* | GRS lab | 2E, 4A |
| MP238 | *CEN1::his3+-PSPOG_00147-mCerulean his3-D1 ura4-D18* | Lorenz lab | 2B-D, 4B, S4A |
| MP316 | *ad6-52 ura4-D18 his3-D1 Padh1-rec8-3HA-ura4+* | This study | 2E |
| MP329 | *ade6-52 ura4-D18 rec11::kanR his3-D1CEN1::his3+-PSPOG_00147-tdTomato* | This study | 4B, S6A-C |
| MP332 | *ade6-52 ura4-D18 rec11::kanR his3-D1 CEN1::his3+-PSPOG_00147-mCerulean* | This study | 4B, S6A-C |
| MP358 | *ade6-52 arg1-14 ura4-D18 rec8::Prec8-Rad21-FLAG-ura4+* | This study | 2E, 3A |
| MP359 | *ade6-M210 ura4-D18 rec8::Prec8-Rad21-FLAG-ura4+* | This study | 2E, 3A |
| MP367 | *CEN1::his3+-PSPOG_00147-tdTomato his3-D1 ura4-D18 Padh1-rec8-3HA-ura4+* | MN lab | 4B, S6A-C |
| MP368 | *CEN1::his3+-PSPOG_00147-mCerulean ura4-D18 his3-D1 Padh1-rec8-3HA-ura4+ C::Padh15-rec11-hygR psc3::KanR* | MN lab | 4B, S6A-C |
| MP370 | *CEN1::his3+-PSPOG_00147-mCerulean his3-D1 ura4-D18 Padh1-rec8-3HA-ura4+* | MN lab | 4B, S6A-C |
| MP381 | *ade6-M210 ura4-D18 rec11::kanR CEN1::his3+-PSPOG_00147-tdTomato his3-D1 rec8D::Prec8-rad21+ -FLAG (GFP?)- ura4+* | This study | 2B-D, S4B-E |
| MP382 | *ade6-M210 ura4-D18 CEN1::his3+-PSPOG_00147-tdTomato his3-D1 rec8D::Prec8-rad21+ -FLAG (GFP?)-ura4+* | This study | 2B-D, 3B, S4B-E |
| MP383 | *ade6-M210 ura4-D18 rec11::kanR CEN1::his3+-PSPOG_00147-mCerulean his3-D1 rec8D::Prec8-rad21+ -FLAG (GFP?)- ura4+* | This study | 2B-D, S4B-E |
| MP385 | *ade6-52 ura4-D18 CEN1::his3+-PSPOG_00147-mCerulean his3-D1 rec8D::Prec8-rad21+ -FLAG (GFP?)-ura4+* | This study | 2B-D, 3B, S4B-E |

| | | | |
|---|---|---|---|
| MP387 | *CEN1::his3+-PSPOG_00147-tdTomato his3-D1 ura4-D18 Padh1-rec8-3HA-ura4+ C::Padh15-rec11-hygR psc3::KanR* | MN lab | 4B, S6A-C |
| MP429 | *CEN1::his3+-PSPOG_00147-tdTomato his3-D1 ura4-D18 Padh1-rec8-3HA-ura4+ C::Padh15-rec11-hygR* | MN lab | 4B, S6A-C |
| MP450 | *ade6-52 ura4-D18 arg1-14 rec11::kanR* | This study | 4A |
| MP451 | *ade6-52 his3-D1 arg1-14 rec11::kanR* | This study | 4A |
| MP453 | *ade6-M210 his3-D1 ura4-D18 rec11::kanR* | This study | 4A |
| MP462 | *CEN1::his3+-PSPOG_00147-tdTomato his3-D1 ura4-D18 C::Padh15-rec11-hygR arg3-D4* | MN lab | 4B, S6A-C |
| MP463 | *CEN1::his3+-PSPOG_00147-mCerulean his3-D1 ura4-D18 C::Padh15-rec11-hygR* | MN lab | 4B, S6A-C |
| MP464 | *CEN1::his3+-PSPOG_00147-mCerulean his3-D1 ura4-D18 Padh1-rec8-3HA-ura4+ C::Padh15-rec11 hygR* | MN lab | 4B, S6A-C |
| MP496 | *ade6-52 arg1-14 ura4-D18 Padh1-rec8-3HA-ura4+* | This study | 2E |
| MP497 | *ade6-M210 ura4-D18 Padh1-rec8-3HA-ura4+* | This study | 2E |
| MP498 | *ade6-52 arg1-1 ura4-D18 c::Padh15-rec11-hygR* | This study | 4A |
| MP499 | *ade6-M210 ura4-D18 c::Padh15-rec11-hygR* | This study | 4A |
| MP500 | *ade6-M210 ura4-D18 c::Padh15-rec11-hygR Padh1-rec8-3HA-ura4+* | This study | 4A |
| MP501 | *ade6-52 arg1-14 ura4-D18 c::Padh15-rec11-hygR Padh1-rec8-3HA-ura4+* | This study | 4A |
| MP502 | *ade6-M210 ura4-D18 psc3:: KanR Padh1-rec8-3HA-ura4+ c::Padh15-rec11-hygR* | This study | 4A |
| MP503 | *ade6-52 ura4-D18 arg1-14 psc3::KanR Padh1-rec8-3HA-ura4+ c::Padh15-rec11-hygR* | This study | 4A |
| MP520 | *ade6-52 rec11-270(5A) arg1-14* | GRS lab | 4A |
| MP521 | *ade6-M26 rec11-270(5A)* | GRS lab | 4A |
| MP545 | *ade6-52 arg1-14 ura4-D18 rec11::kanR rec8D::Prec8-rad21-FLAG-ura4+* | This study | 2E |
| MP546 | *ade6-M210 ura4-D18 rec11::kanR rec8D::Prec8-rad21-FLAG-ura4+* | This study | 2E |
| MP669 | *ura4-D18 his3-D1 rec8::kanR CenI::his3+-PsPOG_00147 tdTomato* | This study | 2B-D, S4A-E |
| MP671 | *ade6-M216 ura4-D18 his3-D1 CenI::his3+-PsPOG_00147 mCerulean rec8::kanR* | This study | 2B-D, S4A-E |
| MP674 | *ade6-M216 ura4-D18 his3-D1 rec8::kanR* | This study | 2E |

| | | | |
|---|---|---|---|
| MP679 | *ade6-52 ura4-D18 his3-D1 rec11::kanR chk1::hygR mid1::his3+* | This study | 4A |
| MP681 | *ade6-52 his3-D1 ura4-D18 arg1-14 rec11::kanR rec8::Prec8-rad21-FLAG-ura4+* | This study | 2E |
| MP682 | *ade6-52 his3-D1 ura4-D18 arg1-14 rec11::kanR rec8::Prec8-rad21-FLAG-ura4+ chk1::hygR mid1::his3+* | This study | 2E |
| MP685 | *ade6-52 his3-D1 ura4-D18 Padh1-rec8-3HA-ura4+ chk1::hygR mid1::his3+* | This study | 2E |
| MP732 | *ade6-52 ura4-D18 his3-D1 rec8::kanR chk1::hygR mid1::his3+* | This study | 2E |
| MP736 | *his3-D1 ura4-D18 rec8::Prec8- rad21-FLAG-ura4+* | This study | 2E |
| MP758 | *ade6-52/M216 ura4+-padh-rec11-HA-kanR padh-rec8-HA-ura4+ ura4-D18 his3-D1* | This study | 4A |
| MP760 | *ade6-M210 ura4-D18 his3-D1 Padh1-rec11-ura4+ psc3::KanR Padh1-rec8-3HA-ura4+ ade6-52 chk1::hygR mid1::his3+* | This study | 4A |
| MP762 | *ade6-M210 ura4-D18 his3-D1 Padh1-rec11-ura4+ psc3::kanR Padh1-rec8-3HA-ura4+* | This study | 4A |
| MP874 | *ura4-D18 rec8::Kan ade6-52 rec11::kanR his3-D1 CEN1::his3+-PSPOG_00147-mCerulean* | This study | 2B-D, S4B-E |
| MP875 | *ade6-52 ura4-D18 rec8::KanR rec11::kanR his3-D1 CEN1::his3+-PSPOG_00147-tdTomato* | This study | 2B-D, S4B-E |
| MP885 | *ade6-52/M216 ura4+-Padh1-rec11-HA-kanR Padh1-rec8-HA-ura+ ura4+-D18 his3-D1 chk1::hygR mid1::his3+* | This study | 4A |
| MP893 | *ura4-D18 his3-D1 Ura4+-Padh1-rad21 CenI::his3+-PsPOG_00147 tdTomato* | This study | 2B-D, S4B-E |
| MP946 | *his3-D1 ura4-D18 rec8::Prec8-rad21-ura4+ chk1::hygR mid1::his* | This study | 2E |
| MP951 | *pat1-114 rec8-3pk-bsdR rec11-HA-kan psc3-Flag-NatR* | This study | 1A-E, 3D, 5A-C, S2B, S2C, S5A, S5B |
| MP969 | *ade6-52 arg1-14 rec11-GFP-rec10-bsdR his3-D1 ura4-D18 rec10::KanR rec8D::Prec8-rad21-FLAG-ura4+* | This study | 3A |
| MP970 | *ade6-52 arg1-14 rec11-GFP-rec10-bsdR ura4-D18 rec10::KanR rec8D::Prec8-rad21-FLAG-ura4+* | This study | 3A |
| MP971 | *ade6-M26 rec11-GFP-rec10-bsdR rec10::KanR ura4-D18 rec8D::Prec8-rad21-FLAG-ura4+* | This study | 3A |
| MP972 | *ade6-M26 rec11-GFP-rec10-bsdR rec10::KanR ura4-D18 his3-D1 rec8D::Prec8-rad21-FLAG-ura4+* | This study | 3A |

| | | | |
|---|---|---|---|
| MP1002 | *pat1-114 rec8::Kan rec11-HA-kanR psc3-Flag-NatR* | This study | 1A-E, 3D, S2C |
| MP1054 | *ade6-M26 ura4-D18 his3-D1 rec11-GFP-rec10-Bsd rec10::KanR CEN1:: his3 PSPOG_00147 mCerulean his3-D1 rec8D::Prec8-rad21-FLAG(GFP?)-ura4+* | This study | 3B |
| MP1055 | *ade6-52 his3-D1 ura4-D18 rec8D::Prec8-rad21-FLAG-ura4+ rec11-272 (5D)* | This study | 3A |
| MP1059 | *ade6-M26 ura4-D18 CEN1::his3+-PSPOG_00147_tdTomato rec8D::Prec8-rad21-FLAG-ura4+ rec10::KanR rec11-GFP- rec10-bsdR his3-D1 ura4-D18* | This study | 3B |
| MP1075 | *rad21-K1::hygR* | NBRP | S5D |
| MP1132 | *ade6-M26/M210 ura4-D18 rec8D::Prec8-rad21-FLAG(GFP?)-ura4+ his3-D1 arg1-14 rec11-272 (5D)* | This study | 3A |
| MP1142 | *ade6-52 pat1-114 rec8-3pk-bsdR rec11::kanR psc3-FLAG-natR* | This study | 5A-C |
| MP1189 | *ura4-D18 his3-D1 ura4+-Padh1-rad21 CEN1::his3+-PSPOG_00147-mCerulean* | This study | 2B-D, S4B-E |
| MP1217 | *ade6-52 pat1-114 rec10-3pk-bsdR rec11::kanR psc3-Flag-natR mug20-myc-hygR* | This study | 5D, 5G |
| MP1218 | *pat1-114 rec10-3pk-bsdR rec11-HA-kanR psc3-Flag-natR mug20-myc-hygR* | This study | 5D, 5G |
| MP1222 | *rec8D::Padh1-rad21-lnk-rec11-HA-kanR ade6-M210 ura4-D18* | This study | 3A |
| MP1248 | *ade6-52 pat1-114 rec10-3pk-bsd rec11::kanR psc3-Flag-natR rec15-myc-hygR* | This study | 5D, 5F |
| MP1249 | *pat1-114 rec10-3pk-bsd rec11-HA-kanR psc3-Flag-natR rec15-myc-hygR* | This study | 5D, 5F |
| MP1363 | *rec8D::Padh1-rad21-lnk-rec11-HA-kanR ade6-52 arg1-14 ura4-D18* | This study | 3A |
| MP1415 | *leu1-32 rad21-GFP-[LEU2] lys1::Padh1-rec11-GBP-mcherry-hygR his3-D1* | This study | 3E |
| MP1450 | *leu1-32 rad21-GFP-[LEU2] lys1::Padh1-rec11-GBP-mcherry-hygR rec8::kanR CEN1::his3+-PSPOG_00147_tdTomato his3-D1* | This study | 3B, 3E |
| MP1451 | *leu1-32 rad21-GFP-[LEU2] lys1::Padh1-rec11-GBP-mcherry-hyg rec8::kanR CEN1:: his3 PSPOG_00147 mCerulean his3-D1* | This study | 3B, 3E |
| MP1473 | *rad21-K1::hygR rec8D::Padh1-rad21-lnk-rec11-HA-KanR* | This study | S5D |

| | | | |
|---|---|---|---|
| MP1499 | *pat1-114 ura4-D18 rec8::kanR ura4+-Padh1-rad21 rec11-HA-kanR psc3-FLAG-natR* | This study | 3C, 3D |
| MP1500 | *leu1-32 rad21-GFP-[LEU2] lys1::Padh1-rec11-GBP-mcherry-hygR his3-D1* | This study | 3E |

**Table S2** - List of oligonucleotides used in the study.

| Oligomer # | Sequence | Purpose |
|---|---|---|
| MO8 | CGATAGTGGAAACCGACG | *3xMyc-hygR* integration check into *mug20* |
| MO90 | GAATCATTCCATCCACTTCTTTCTC | *ura4+*- integration check into *rad21* |
| MO100 | CTGTCTTCGGTATCGTCGTATC | *3xMyc-hygR* integration check into *mug20* |
| MO113 | CGCAAGGAATCGGTCAATAC | *3xMyc-hygR* integration check into *rec15* |
| MO136 | GGAGAGCAGTAGCTGGTTCA | *rad21* homology region amplification *and ura4+-Padh1* integration check into *rad21* |
| MO215 | GGAGATTCATCCTTCAATTCTC | *rad21* homology region amplification |
| MO216 | ATACTGTAAGGTTTGCAATTCTGTTGGAAGAAGGCAATTC | *rad21* homology region amplification |
| MO217 | AATTGCAAACCTTACAGTATGGAATGCCATGTCAGATTTG | *ura4+-Padh1* amplification |
| MO218 | AGCTGATCTCAGATCTGGGGATCCGCTAGCGGAATTCT | *ura4+-Padh1* amplification |
| MO219 | CCAGATCTGAGATCAGCTATGTTCTATTCAGAGGCCATTC | *rad21* homology region amplification |
| MO415 | AGCGAGTACTGGCTATTGATTTCTTAGCAGATGACGATTA TATTCTGGCGGGGATAATT | *rec15-3xMyc-HygR* generation |
| MO416 | AAAATCAAACTATGCCCAGAAGTAATACAACAACGAAAAA GCGAGTACTGGCTATTGAT | *rec15-3xMyc-HygR* generation |
| MO417 | TCATAGAAAAGCTTTTAAATGAGGCTGATCTAAGTATTTAG TAACTAATGCATTCGTGA | *rec15-3xMyc-HygR* generation |
| MO418 | AAAACGGTTTGATTGGTGTTTAATTTTAATTGATATCATAG AAAAGCTTTTAAATGAGG | *rec15-3xMyc-HygR* generation |
| MO419 | TGATTGAAACCTCAACTCATAAAGCTATTCTCGACAATTTT ATTCTGGCGGGGATAATT | *mug20-3xMyc-HygR* generation |
| MO420 | CATATTACTAACCTCAATAAATTTAAAGAAACTAAAGATGA TTGAAACCTCAACTCATA | *mug20-3xMyc-HygR* generation |
| MO421 | GCTTGGTTTGAGGCAGTATGACATTAGATCTCTGAAATTA GTAACTAATGCATTCGTGA | *mug20-3xMyc-HygR* generation |
| MO422 | AGCGTGTTTGATGAAAGTTGACTTTCTTCGTTATTATCAAG CTTGGTTTGAGGCAGTAT | *mug20-3xMyc-HygR* generation |

| | | |
|---|---|---|
| MO430 | GGCTTTCATAAGTCTATCAACAG | *3xMyc-hygR* integration check into *rec15* |
| MO439 | AAAGAAAGCGGCAGCGTGAGCAGCGAACAGCTGGCGCAGTTTCGCAGCCTGGATGAAGGC | *rad21-lnk-rec11-HA-kanR* generation |
| MO440 | CGCAGCCTGGATGAAGGCAAAAGCAGCGGCAGCGGCAGCATGAGATTCGAATTTGAATCA | *rad21-lnk-rec11-HA-kanR* generation |
| MO441 | TTATATCTGTCAAGCAGGATGTTGCCATTCAGAACGAAATCACGCTTACTGCTAAACGTGGAATGCTACTTTCATCACTAAAAGAAAGCGGCAGCGTGAG | *rad21-lnk-rec11-HA-kanR* generation |
| MO442 | ATTATTTTGACAACTTCAACAAAGGGTTTAATTCCCAAATTCAACGTTTTAAAAATTTTAAGTTTAACATGTGAAAAGTTGAATTCGAGCTCGTTTAAAC | *rad21-lnk-rec11-HA-kanR* generation |
| MO493 | AGTCAGTGGGTACCATGAGATTCGAATTTGAATC | *rec11-GBP-mcherry* generation |
| MO494 | GACTGACTGGATCCTGATAAATGGTCAGCTGTTT | *rec11-GBP-mcherry* generation |
| MCO1 | CAACTTACATCAGCATACTGG | ChIP qPCR- *cnt* all |
| MCO2 | CGAAAGATGGTCAATTGCTT | ChIP qPCR- *cnt* all |
| MCO5 | AAAGCAAAAACCGAGTTTGG | ChIP qPCR- *imrIII* |
| MCO6 | AAATTTCATGAACGCTGACG | ChIP qPCR- *imrIII* |
| MCO15 | TGTGCCTCGTCAAATTATCATCCATCC | ChIP qPCR- *dg* all |
| MCO16 | ACTTGGAATCGAATTGAGAACTTGTTATGC | ChIP qPCR- *dg* all |
| MCO25 | GGAATAGCATACCGTCAAGTCGTTAGTTG | ChIP qPCR- *dh* all |
| MCO26 | GGTCAACGCACGCCTAAACTAGC | ChIP qPCR- *dh* all |
| MCO33 | GAAATCTAGTCGAGGTCAAG | ChIP qPCR- *msp1* |
| MCO34 | CTTCCAAGTACTGCAAAACC | ChIP qPCR- *msp1* |
| MCO37 | CGTCCAATTTAGGCCAT | ChIP qPCR- *bub1* |
| MCO38 | GTCTTAACCGTCTCCCTG | ChIP qPCR- *bub1* |
| MCO39 | TGGCTCTAGATGTATCAGCG | ChIP qPCR- *sib1* |
| MCO40 | ACGGCTGTCGAATCAGTTAC | ChIP qPCR- *sib1* |
| MCO41 | AAACACTGGTCATCTCGGAACCAGC | ChIP qPCR- *mbs1* |
| MCO42 | TCCTTTGCAGTGAATGGTTGGGCTAC | ChIP qPCR- *mbs1* |
| MCO43 | TCATCTCATTAAGAGTGTTCGG | ChIP qPCR- *sat1* |
| MCO44 | GATGCAAAGCAAATGCGCAAGT | ChIP qPCR- *sat1* |
| MCO45 | TGCCATGAATTCCTATCGTAG | ChIP qPCR- *SPBC3H7.03* |

| | | |
|---|---|---|
| MCO46 | GTTGCAAGGCTTATGTACATAC | ChIP qPCR- *SPBC3H7.03* |
| MCO47 | CTTGCTCTGTAGCTTAGGAG | ChIP qPCR- *birI* |
| MCO48 | GACAGAAACGTCGGAAACTC | ChIP qPCR- *birI* |

**Table S3.** Meiotic segregation defects at *cen1* to determine complementation by Rad21.

| S. No. | Genotype | Strains crossed | Trials | Total tested | Segregation pattern (# of tetrads) | | | | | | | |
|---|---|---|---|---|---|---|---|---|---|---|---|---|
| | | | | | Normal | RS | | Meiosis I errors | | | | Meiosis II errors |
| | | | | | | | | NDJ | | PSSC | | |
| | | | | | | RS | RS+MII error | NDJ | NDJ+MII error | PSSC | PSSC+MII error | |
| 1 | *Wild type* | MP208 X MP238 | 1 | 134 | 131 | 2 | 0 | 0 | 0 | 0 | 0 | 1 |
| | | | 2 | 177 | 171 | 5 | 0 | 0 | 0 | 0 | 0 | 1 |
| | | | 3 | 90 | 88 | 0 | 0 | 0 | 0 | 0 | 0 | 2 |
| | | | 4 | 162 | 153 | 5 | 1 | 0 | 1 | 1 | 0 | 1 |
| 2 | *rad21oe* | MP893 X MP1189 | 1 | 205 | 191 | 3 | 3 | 0 | 0 | 0 | 1 | 7 |
| | | | 2 | 176 | 160 | 6 | 2 | 0 | 0 | 0 | 0 | 8 |
| | | | 3 | 125 | 114 | 6 | 0 | 0 | 0 | 0 | 0 | 5 |
| 3 | *rec8Δ* | MP669 XMP671 | 1 | 166 | 0 | 23 | 40 | 0 | 18 | 0 | 79 | 6 |
| | | | 2 | 105 | 0 | 7 | 26 | 0 | 18 | 5 | 42 | 7 |
| | | | 3 | 91 | 0 | 6 | 21 | 0 | 27 | 3 | 30 | 4 |
| 4 | *rec8Δ rec11Δ* | MP874 X MP875 | 1 | 153 | 0 | 8 | 44 | 0 | 48 | 1 | 38 | 14 |
| | | | 2 | 55 | 0 | 6 | 15 | 0 | 12 | 1 | 19 | 2 |
| 5 | *rec8Δ rad21ee* | MP382 X MP385 | 1 | 100 | 0 | 8 | 19 | 0 | 27 | 0 | 41 | 5 |
| | | | 2 | 59 | 0 | 1 | 15 | 0 | 16 | 1 | 22 | 4 |
| | | | 3 | 279 | 0 | 31 | 70 | 0 | 56 | 3 | 106 | 13 |
| 6 | *rec8Δ rec11Δ rad21ee* | MP381 X MP383 | 1 | 92 | 0 | 10 | 21 | 0 | 23 | 0 | 35 | 3 |
| | | | 2 | 124 | 0 | 17 | 34 | 2 | 19 | 3 | 45 | 4 |
| | | | 3 | 179 | 0 | 19 | 54 | 0 | 36 | 0 | 63 | 7 |
| 7 | *rec8Δ rad21ee rec11-gfp-rec10 rec10Δ* | MP1054 X 1059 | 1 | 124 | 0 | 14 | 26 | 0 | 29 | 0 | 45 | 10 |
| | | | 2 | 101 | 0 | 7 | 24 | 0 | 21 | 0 | 47 | 2 |
| | | | 3 | 70 | 0 | 6 | 16 | 0 | 14 | 0 | 32 | 2 |
| | | | 4 | 78 | 0 | 8 | 23 | 1 | 15 | 0 | 25 | 6 |
| 8 | *rec8Δ rad21-gfp rec11-gbp* | MP1450 X MP1451 | 1 | 110 | 0 | 13 | 25 | 0 | 20 | 0 | 48 | 4 |
| | | | 2 | 102 | 0 | 10 | 30 | 0 | 15 | 1 | 40 | 6 |
| | | | 3 | 104 | 0 | 9 | 28 | 1 | 19 | 1 | 45 | 1 |

**Table S4.** Meiotic segregation defects at *cen1* to determine complementation by Psc3.

| **Sr. No.** | **Genotype** | **Strains crossed** | **Trials** | **Total tested** | **Segregation pattern (# of tetrads)** | | | | | | | |
|---|---|---|---|---|---|---|---|---|---|---|---|---|
| | | | | | Normal | RS | | Meiosis I errors | | | | Meiosis II errors |
| | | | | | | | | NDJ | | PSSC | | |
| | | | | | | RS | RS+MII error | NDJ | NDJ+MII error | PSSC | PSSC+MII error | |
| 1 | *rec8oe* | MP367 X MP370 | 1 | 163 | 150 | 3 | 0 | 0 | 0 | 2 | 0 | 8 |
| | | | 2 | 217 | 197 | 1 | 1 | 1 | 1 | 3 | 0 | 13 |
| | | | 3 | 79 | 66 | 0 | 0 | 0 | 1 | 4 | 2 | 6 |
| 2 | *rec11oe* | MP462 X MP463 | 1 | 186 | 176 | 3 | 0 | 0 | 0 | 0 | 2 | 5 |
| | | | 2 | 210 | 199 | 3 | 5 | 0 | 0 | 0 | 0 | 3 |
| | | | 3 | 263 | 253 | 5 | 2 | 0 | 1 | 0 | 0 | 2 |
| 3 | *rec8oe rec11oe* | MP429 x MP464 | 1 | 143 | 126 | 3 | 3 | 0 | 1 | 0 | 1 | 9 |
| | | | 2 | 206 | 192 | 4 | 3 | 0 | 0 | 0 | 0 | 7 |
| | | | 3 | 111 | 104 | 3 | 0 | 0 | 0 | 0 | 0 | 4 |
| 4 | *rec11Δ* | MP329 X MP332 | 1 | 129 | 50 | 4 | 2 | 13 | 11 | 27 | 2 | 20 |
| | | | 2 | 152 | 80 | 3 | 3 | 21 | 4 | 18 | 3 | 20 |
| | | | 3 | 111 | 56 | 0 | 4 | 6 | 13 | 12 | 4 | 16 |
| | | | 4 | 137 | 76 | 2 | 1 | 19 | 4 | 15 | 4 | 16 |
| | | | 5 | 85 | 44 | 0 | 1 | 9 | 6 | 15 | 1 | 9 |
| 5 | *psc3Δ rec8oe rec11oe* | MP368 X MP387 | 1 | 76 | 13 | 2 | 3 | 0 | 4 | 4 | 3 | 47 |
| | | | 2 | 87 | 19 | 2 | 3 | 0 | 1 | 5 | 14 | 43 |
| | | | 3 | 126 | 37 | 6 | 6 | 0 | 1 | 1 | 11 | 64 |

**Table S5.** Arm recombination between *ade6-arg1* interval during meiosis.

| **S. No.** | **Genotype** | **Strains crossed** | **No. of crosses** | **Total tested** | **Arm recombination (*ade6-arg1*)** | |
|---|---|---|---|---|---|---|
| | | | | | No. of recombinants | Recombinant Frequency (RF; %) |
| 1 | *Wild type* | MP4 X MP96 | 4 | 509 | 160 | 31.55 |
| 2 | *rec8oe* | MP496 X MP497 | 3 | 396 | 131 | 33 |
| 3 | *rec8Δ rad21ee* | MP358 X MP359 | 4 | 527 | 4 | 0.775 |
| 4 | *rec8Δ rec11Δ rad21ee* | MP545 X MP546 | 3 | 396 | 1 | 0.25 |
| 5 | *rec8Δ rad21ee rec11-5D* | MP1132 X MP1055 | 4 | 406 | 3 | 0.66 |
| 6 | *rec8Δ rad21ee rec11-rec10* | MP969/970 X MP971/972 | 3 | 567 | 48 | 8.33 |
| 7 | *rec8Δ rad21-rec11oe* | MP1222 X MP1363 | 3 | 393 | 1 | 0.25 |
| 8 | *rec11oe* | MP498 X MP499 | 3 | 396 | 134 | 33.82 |
| 9 | *rec8oe rec11oe* | MP500 X MP501 | 3 | 396 | 131 | 33.07 |
| 10 | *rec11Δ* | MP450/451 X MP453 | 3 | 400 | 5 | 1.27 |
| 11 | *rec11(5A)* | MP520 X 521 | 3 | 386 | 4 | 1.07 |
| 12 | *psc3Δ rec8oe rec11oe* | MP502 X MP503 | 3 | 396 | 103 | 25.97 |

**Table S6.** Centromeric recombination at *chk1-mid1* interval during meiosis.

| **Sr. No.** | **Genotype** | **Strains crossed** | **No. of crosses** | **Total tested** | **Cen recombination (*chk1-mid1*)** | |
|---|---|---|---|---|---|---|
| | | | | | No. of recombinants | Recombinant Frequency (RF; %) |
| 1 | *Wild type* | MP172 x MP209 | 3 | 395 | 0 | 0 |
| 2 | *rec8oe* | MP685 x MP316 | 3 | 396 | 0 | 0 |
| 3 | *rec8Δ* | MP674 x MP732 | 3 | 371 | 1 | 0.27 |
| 4 | *rec8Δ rad21ee* | MP946 x MP736 | 3 | 396 | 0 | 0 |
| 5 | *rec8Δ rec11Δ rad21ee* | MP681 x MP682 | 3 | 395 | 0 | 0 |
| 6 | *rec8oe rec11oe* | MP885 x MP758 | 3 | 369 | 0 | 0 |
| 7 | *rec11Δ* | MP679 x MP41 | 3 | 387 | 0 | 0 |
| 8 | *psc3Δ rec8oe rec11oe* | MP760 x MP762 | 3 | 396 | 0 | 0 |
